# Structural promiscuity in the human circulatory IgA1 clonal repertoire

**DOI:** 10.1101/2025.11.13.688237

**Authors:** Amber D. Rolland, Sofia Kalaidopoulou Nteak, Gestur Vidarsson, Albert Bondt, Albert J. R. Heck

## Abstract

Immunoglobulin A (IgA) is the most abundant antibody in humans, with high concentrations in both mucosae/secretions and circulation. While mucosal IgA has been studied extensively, characterization of human IgA in serum and its distinctive functions lags behind. Circulatory IgA is regularly assumed to be monomeric, despite some reports describing a minor population of J-chain-coupled dimers. Here, we first charted the compositional landscape of human serum IgA in individual healthy donors. In addition to the expected predominating monomers, we consistently observed J-coupled dimers, even representing ∼30% of one donor’s total serum IgA. To determine whether these structurally distinct populations were also clonally distinct, we employed mass spectrometry-based IgA1 clonal profiling in sera of two donors. Our data revealed the majority of IgA1 clones are present solely as monomers, with a smaller number exclusively dimeric. Strikingly, a third population of IgA1 clones is present in circulation as both monomers and J-coupled dimers. In fact, these shared, structurally-promiscuous IgA1 clones dominated both individuals’ serum IgA1 clonal repertoires. Our findings suggest likely every J+ IgA-secreting cell can co-produce monomers and J-coupled dimers, but what exactly determines this ratio requires further investigation. This finding is important, as IgA monomers and J-coupled dimers have distinct characteristics in antigen binding, receptor activation, and clearance from circulation, with several reports highlighting, for instance, the enhanced neutralization capacity of dimeric IgA. As we show here, the human immune system is not merely capable of producing both forms, but apparently prefers doing so in parallel.

**Significance Statement:** Human immunoglobulin IgA occurs in diverse assemblies, mainly monomers (mIgA) and J-chain coupled dimers (dIgA). The general view is that these forms are produced at different sites, i.e., circulatory and mucosal, with the former assumed to represent mIgA. Inherently, it would not be expected that these populations have any overlap in their clonal repertoire. Here, we report the first analysis at the protein level of assembly-specific IgA1 clonal repertoires in sera of healthy individual donors. Strikingly, we find a substantial subset of clones that co-occur as both mIgA1 and dIgA1 assemblies, hinting at shared B-cell origins. This marks a paradigm shift in current understanding of human IgA, where these assemblies are classically viewed as structurally, functionally, and clonally distinct.

## Introduction

Of all classes of immunoglobulins in humans, immunoglobulin A (IgA) is the predominant isotype found in the mucosal linings of the gastrointestinal, respiratory, and urogenital tracts and in external secretions such as tears, saliva, and colostrum/milk (***1, 2***). IgA is also prevalent in the lymphatic and circulatory systems, ranking as the second-most abundant Ig in adult serum (∼0.5-3.5 mg/mL) after IgG (***1***). Together these high concentrations in both mucosal and systemic immune compartments make IgA the overall most abundant Ig in the human body (***3***).

All human Ig classes—IgM, IgD, IgE, IgG, IgA—have relatively similar domain structure due to their formation via class-switch recombination. They share the same basic protomer (H_2_L_2_) structure of two heavy (H) chains paired with two light (L) chains expressed and assembled by antibody-secreting cells (ASCs). IgG is present exclusively in this monomeric form in both serum and mucosal-associated biofluids/tissues. By contrast, human IgA is largely present as polymers in mucosal contexts. The presence of a C-terminal 18-residue peptide extension called the tailpiece (TP) in both IgA and IgM enables formation of J-coupled polymeric IgA or IgM (pIgA, pIgM) assemblies (***4***). This process depends upon incorporation of a single molecule of the immunoglobulin joining chain, or J-chain (J), inside the ASC before secretion (***5, 6***). While IgM exclusively assembles into J-containing pentamers which associate extracellularly with CD5L in the sera of healthy individuals (***7***), this degree of structural specificity induced by the J-chain is not mirrored in IgA. *In vitro* studies have demonstrated as many as five IgA protomers can be assembled alongside a single J-chain (***8***). However, such large structures are typically only found in trace amounts in humans—if detected at all (***1***). Instead, pIgA is by far predominantly found in the form of J-coupled dimers (dIgA) (***2***), which will be the central focus of the present work (Fig. 1A). Regardless of the stoichiometry, the presence of the J-chain in polymeric assemblies of IgA or IgM enables binding to the polymeric immunoglobulin receptor (pIgR) essential for transepithelial transport into the mucosae (***9***). There, proteolytic cleavage of pIgR releases the J-coupled dIgA into the mucosal lumen with part of pIgR’s N-terminal extracellular domain—called the secretory component (SC)—still attached. Together, the pIgA-J-SC complex constitutes a distinct molecular form (secretory IgA, SIgA) which is specific to mucosal contexts (Fig. 1A). There, it forms the first line of defense in protecting the intestinal epithelial barrier from enteric infection and invasion by pathogenic microorganisms (***1***).

**Figure 1.**
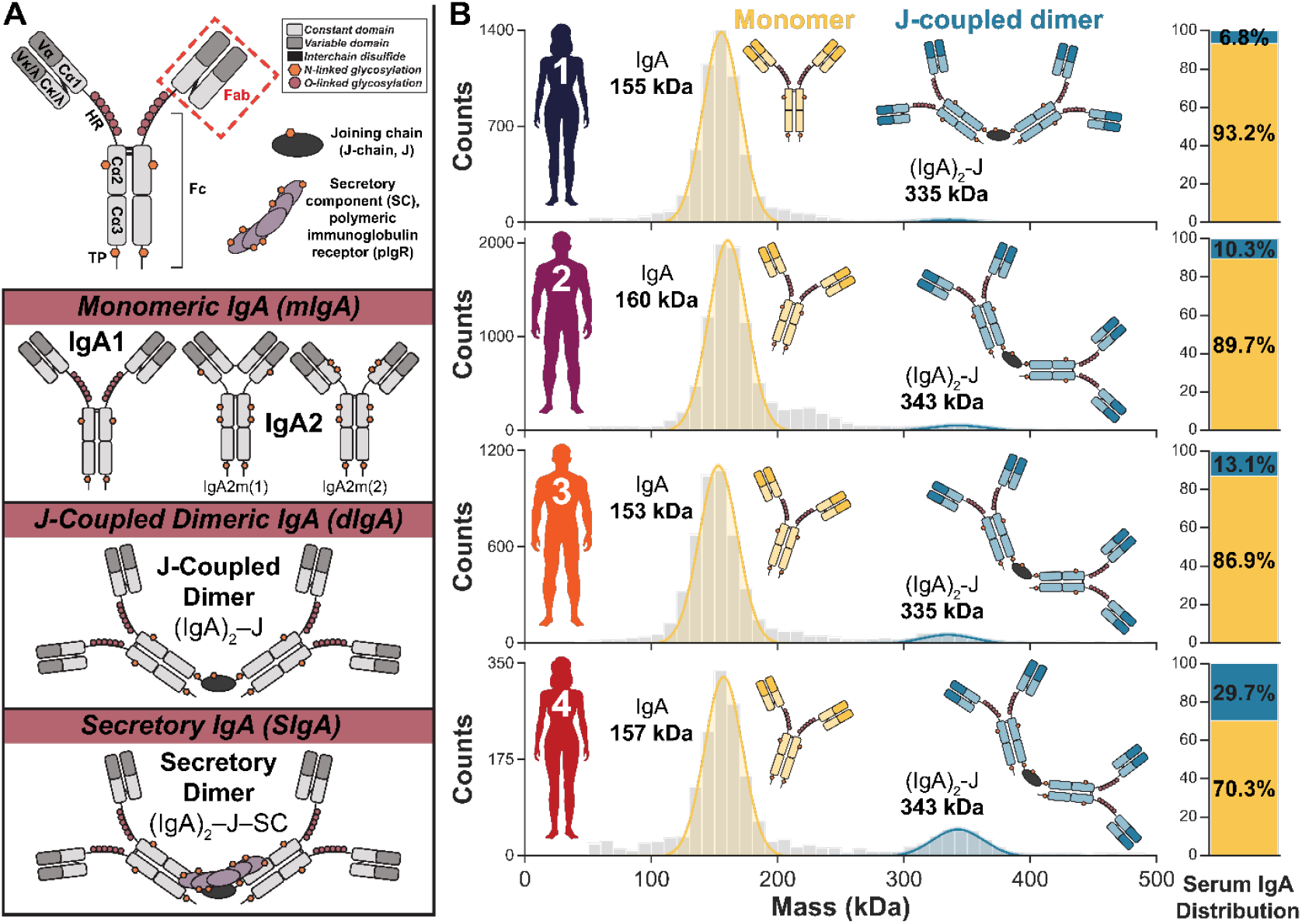
Human serum can contain a substantial amount of J-coupled dimers of IgA. **(A)** Schematic diagrams of human IgA1 monomer **(top)** with domains, disulfide bonds, and glycosylation sites highlighted, along with the J-chain (J) and secretory component (SC) of pIgR assembly factors. Below are schematic models illustrating the molecular heterogeneity and structural variability of human IgA. The three distinct molecular forms are defined by the absence of both assembly factors (monomeric IgA, mIgA), presence of only J-chain without SC of pIgR (dimeric IgA, dIgA), and presence of both the J-chain and SC of pIgR (secretory IgA, SIgA) as indicated. Schematic models of monomeric human IgA1, IgA2m(1), and IgA2m(2) highlight the differences in structural features between the two subclasses and allotypes. **(B)** Mass distributions of the full repertoire of intact IgA assemblies purified from serum of four adult donors (Donors 1-4 as indicated) determined by mass photometry. Two assemblies were detected in all four samples and assigned as monomers (yellow, ∼160 kDa) and J-coupled dimers (blue, ∼340 kDa). These assignments were verified by mass-spectrometry based bottom-up proteomics. The relative abundances of these assemblies varied between individuals, as indicated by the bar charts to the right of each panel which depict the percentage of total circulatory IgA present as J-coupled dimers (blue) or monomers (yellow). The total number of individual monomer and J-coupled dimer molecules detected (represented by the color-shaded area of each assigned, fitted peak) were quantified and then adjusted accordingly to reflect the presence of two IgA monomer subunits in each J-coupled dimer.

Incorporation of the J-chain thus gives rise to aspects of human IgA’s uniquely extensive molecular heterogeneity (***10***). Moreover, J-chain association is merely optional rather than required for production of human IgA, which consequently exhibits structural promiscuity whereby both monomeric IgA lacking the J-chain (mIgA) and J-coupled polymeric forms (including dIgA) naturally occur, a uniquely distinct feature compared to all other human Ig isotypes. With two subclasses (namely IgA1 and IgA2, Fig. 1A) found only in humans and some other great ape species, the human IgA system is also distinct compared to most other mammalian IgA (***11***). Human IgA1 and IgA2 are highly similar with ≥90% sequence homology between the three Cα domains (***2***). Differences are almost exclusively restricted to the hinge region (HR) which is 13 amino acids longer in IgA1, featuring two Pro/Ser/Thr-rich repeats that introduce numerous putative O-glycosylation sites exclusively to this subclass (Fig. 1A). While the extended IgA1 hinge confers greater conformational flexibility, it also makes IgA1 susceptible to cleavage by specific bacterial proteases. Further structural and functional differences between IgA subclasses arise due to their distinct glycosylation profiles (***12-15***). The distribution of subclasses as well as molecular composition varies throughout the body (***16***).

Given the dominance of IgA at mucosal surfaces and in secretions, human IgA has been extensively characterized in the context of mucosal immunology (i.e., SIgA) (***8, 9, 17***). The functions of human IgA in serum are less well characterized. While the structural and molecular composition is clearly distinct from that of mucosal IgA, several basic details still have not been exactly defined in serum (***18, 19***), and such characterization is especially lacking in the context of healthy individuals. Serum IgA is dominated by the IgA1 subclass and is often assumed (or implied) to be exclusively monomeric and thus devoid of the J-chain. However, a few studies have reported the presence of a second, minor population corresponding to J-coupled dimers, but with discrepancies in the reported fractional abundances which range from negligible (<1%) up to ∼22% of total serum IgA (***10, 20, 21***). Whether such values refer to bulk population measurement or were derived from observations within a single donor is often unclear. With nearly no pIgR present in serum, these J-coupled dimers of IgA in circulation are devoid of this co-factor and thus distinct from SIgA.

Serum IgA, like any Ig, in fact represents a molecular mixture of thousands to millions of different IgA1 clonal variants (***22***). Each of these has a unique amino acid sequence in the variable regions contained within the Fab arms that dictates antigen specificity and reflects their distinct cellular origins (***23***). We previously found that endogenous IgG1 and IgA1 repertoires are often dominated by just hundreds of unique clones (***24-26***). Whether these dominant IgA1 clones correspond to mIgA1 or dIgA1 is completely unknown but critical to fully understand human IgA. Not only do monomers and J-coupled dimers exhibit structural and functional differences, but they have also been suggested to originate from distinct IgA+ ASC populations located in distinct immune compartments (***27, 28***).

Here we aim to decipher the exact molecular and clonal composition of human serum IgA1. Using single-molecule mass photometry and mass spectrometry-based proteomics to identify IgA assemblies purified from sera of four healthy individual donors, we chart the varied compositional landscape of human serum IgA. For two of these healthy donors, we further utilize a mass spectrometry-based approach to monitor endogenous IgA1 Fab molecules from each assembly population (mIgA versus dIgA) separately. We report the assembly composition of human serum IgA1 with clonal resolution, annotating the intact molecular form(s) for nearly 3,000 distinct endogenous IgA1 clones. Importantly, we identify novel features of IgA’s inherent structural promiscuity related to J-chain incorporation. These findings have significant implications for understanding the cellular origins, functions, and complexity of human circulatory IgA.

## Results

### In healthy individuals up to one-third of serum IgA can be present as J-coupled dimers

The first unresolved question we aimed to address was whether human circulatory IgA exists exclusively as monomers or exhibits widespread structural promiscuity. All possible molecular forms of circulatory IgA— i.e., mIgA and pIgA assemblies lacking SC—can readily be discriminated by mass and the defining absence or presence of J-chain, respectively (Fig. 1A). We therefore investigated the molecular composition of intact IgA assemblies, affinity-purified from the serum of four healthy adult donors (Donors 1-4, SI Table S1) by using single-molecule mass photometry (MP). The resulting MP mass histograms (Fig. 1B) depict the distribution of all detected particles, allowing identification of each species by mass and quantitative comparison between co-occurring populations.

The distribution of circulatory IgA assemblies from each of the four healthy donors featured as expected a dominant peak at ∼160 kDa, assigned as IgA monomers (Fig. 1B). We also detected the presence of J-coupled dimers at ∼340 kDa for all four donors. We highlight the absence of larger assemblies in circulation, as evidenced by the lack of any peaks consistent with the associated theoretical masses (***26***). These assignments are supported by bottom-up mass spectrometry (MS)-based proteomics analysis of these samples (SI Fig. S1A).

As expected, J-coupled dimers were less abundant than monomers in all four donors (Fig. 1B). However, the intact assembly distribution varied considerably, with roughly 7%, 10%, 13%, and 30% of the total IgA of Donors 1-4, respectively, present as J-coupled dimers without SC of pIgR. For this quantitative analysis we adjusted the particle counts considering that each dIgA molecule detected contains two copies of IgA monomer subunits (SI Table S2 and Supplementary Materials and Methods). The assembly distributions of human IgA in serum reported in the literature vary substantially, with J-coupled dimers accounting for anywhere from as low as <1% up to as high as ∼22% of all circulatory IgA (***20, 21***). However, the consensus view seems to be that there is no (or effectively zero) dIgA in circulation. Although we analyzed only four donor samples here, we observed a substantial amount of dIgA in each, indicating that J-coupled dimers are more prevalent in the circulation of healthy individuals than previously thought.

To further contextualize results from these four donors, we analyzed a publicly available plasma proteomics dataset (***29***) of 687 samples from 139 individuals for the abundances of IgA, IgM, J-chain, and CD5L. Assuming that all J-chain protein detected in circulation—in accordance with the known requirements for proper folding and secretion (***5, 6, 30***)—must be in complex either with IgM pentamers associated with CD5L (IgM_5_-J-CD5L) or with dIgA (IgA_2_-J) (***7, 31***), the estimated relative abundance of dIgA can be derived from the MaxLFQ values of these four proteins. Using this approach described in greater detail in the Supplementary Materials and Methods, we observed that the fractional abundance of J-coupled dimers also varies extensively between the hundreds of individual donors in this dataset (***29***), which indeed ranges from as low as <1% up to as high as ∼33% of the total serum IgA (SI Fig. S1C), validating our data.

### Monomers and J-coupled dimers of IgA1 are clonally diverse in human serum

Our next aim was to dissect endogenous circulatory IgA1 at the protein level with clonal resolution to determine whether the monomeric and J-coupled dimeric repertoires have shared clonality or are mutually exclusive. We previously established methods for profiling endogenous IgA1 Fabs in serum as well as in breastmilk but did not probe their precise molecular composition in these studies (***24-26***). Here, we adapted this earlier approach for the assembly-specific characterization of endogenous circulatory IgA1 clonal repertoires (Fig. 2A and SI Fig. S2).

**Figure 2.**
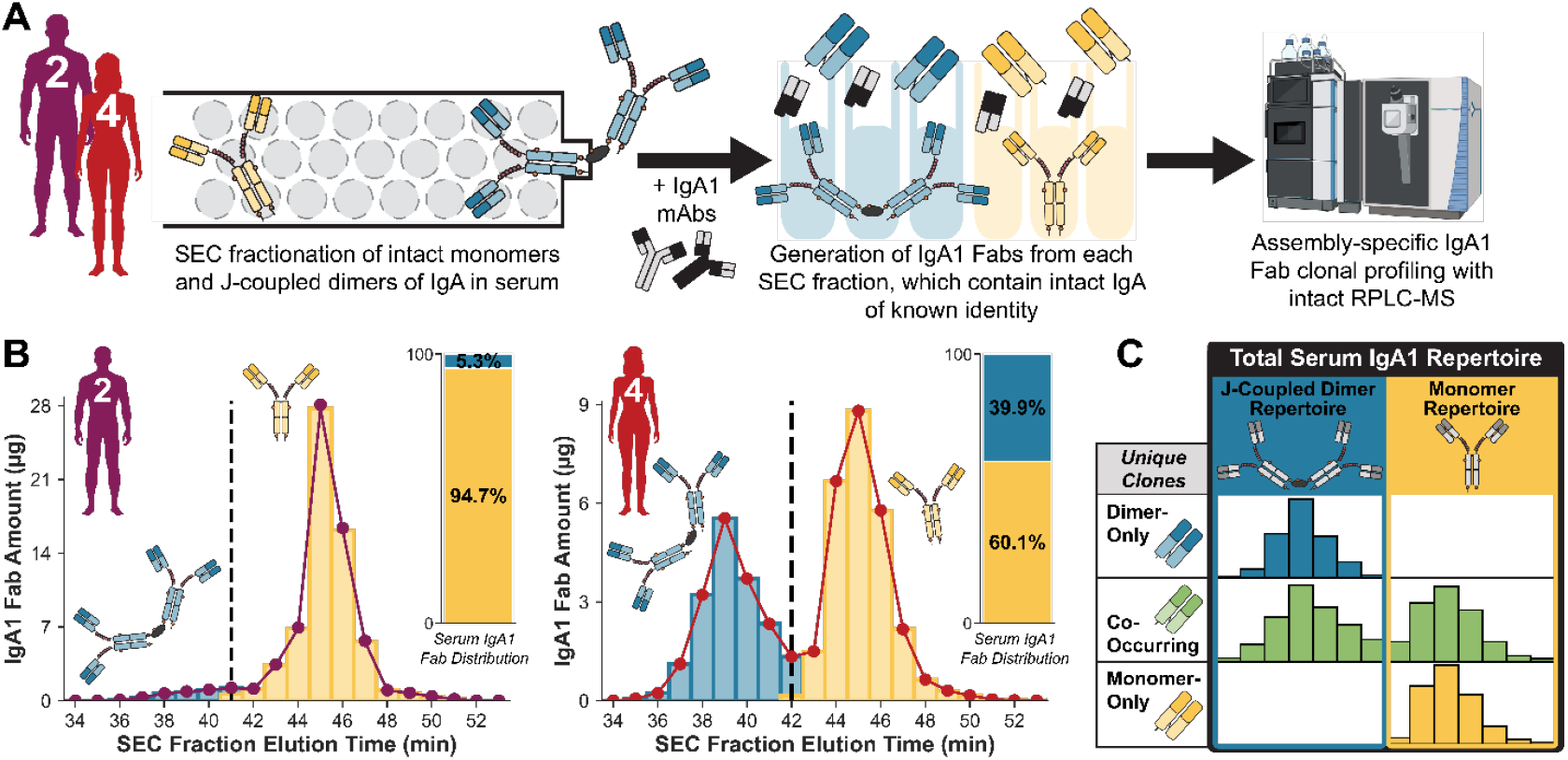
Assembly-specific profiling of IgA1 clonal repertoires reveals structural promiscuity of individual clones. **(A)** Approach taken for the profiling of IgA1 Fab clonal repertoires. Size exclusion chromatography was used to separate intact monomer and J-coupled dimers present in serum, followed by generation of IgA1 Fabs from each monomer- or dimer-containing SEC fraction individually. After spiking in two recombinant monomeric IgA1 mAbs into each SEC fraction to enable intensity-based quantitation, all IgA was captured with an affinity resin, which was then washed to remove unbound non-IgA proteins. Digestion with an O-glycan-specific protease resulted in the release of IgA1 Fabs from each SEC fraction, which were subsequently analyzed by intact RPLC-MS. **(B)** Cumulative IgA1 Fab abundance detected and quantified by intact RPLC-MS in each SEC fraction sample, effectively reconstructing the SEC elution profiles of the intact circulatory IgA1 assemblies for Donor 2 **(left)** and Donor 4 **(right)**. Peaks are annotated as monomers (yellow) or J-coupled dimers (blue), and the inset bar chart depicts the cumulative abundance of all detected IgA1 Fabs from each of the two assemblies. **(C)** Visual reference for key concepts and terminology used. Most unique Fab clones originated exclusively from monomers (“monomer-only”, yellow), with a smaller but appreciable number exclusively found as J-coupled dimers (“dimer-only”, blue). An intriguing third subset of unique IgA1 clones co-occurred in circulation in both intact monomer and J-coupled dimer forms simultaneously (“co-occurring”, green). These three clone populations comprise the “total” serum IgA1 repertoire (black box) which can also be divided into two assembly-specific “J-coupled dimer” (blue box) and “monomer” (yellow box) repertoires as indicated in the illustration.

The circulatory IgA of Donors 1-4 were all similarly dominated by the IgA1 subclass (≥85%) based on MS-based proteomics and therefore suitable for IgA1-based repertoire analysis (SI Fig. S1B). From the four individuals, we selected Donors 2 and 4, who had a normal (∼10%) and high (∼30%) proportion of dIgA, respectively (Fig. 1B), for more detailed clonal analysis. We used offline size exclusion chromatography (SEC) to separate intact monomeric and J-coupled dimeric IgA assemblies in the serum of Donors 2 and 4 (Fig. 2A). Each donor’s intact J-coupled dimers and monomers of IgA were collected in discrete sets of fractions (Fig. 2A). Single-molecule MP analysis of SEC-fractionated affinity-purified IgA provided mass-based evidence for the elution range of each intact assembly (SI Fig. S2A), and MS-based proteomic analysis of all fractions confirmed the range over which IgA and J-chain co-elute as J-coupled dimers (SI Fig. S2B and C).

With the intact form of IgA present in each SEC fraction thus clearly defined based on elution time, we generated IgA1 Fab molecules for each IgA-containing SEC fraction individually (Fig. 2A) as detailed further in the Supplementary Materials and Methods (***24, 26***). Using this approach, distinct IgA1 Fab clones are identified by their characteristic unique mass and reversed phase liquid chromatography (RPLC) retention time (RT), and the abundance of each of these unique clones in each fraction is determined. This allows reconstruction of the intact IgA1 SEC elution profile—using the cumulative detected abundance as in Fig. 2B or that of a single distinct clone—and assignment of assembly state, i.e., mIgA1 or dIgA1 (Fig. 2B). As expected, the cumulative fractional abundances of Fabs originating from intact IgA1 monomers or separately from J-coupled dimers (Fig. 2B) differed considerably between Donors 2 and 4, in line with their intact assembly distributions observed by MP (Fig. 1B). Also as anticipated, the unique IgA1 clones present as monomers outnumbered the unique J-coupled dimer clones, and this difference was markedly higher for Donor 2 than for Donor 4 (SI Fig. S3). Importantly, we found both the monomer and J-coupled dimer populations to be unique and clonally diverse, with hundreds of distinct IgA1 clones identified for each assembly state in both donors (SI Fig. S3A and B). Further, with respect to Fab mass and RPLC RT, we observed similar distributions between both assembly-specific clonal populations—i.e., the variable Fab regions from mIgA1 versus dIgA1 antibodies do not fall within obviously distinct mass windows or RT ranges (SI Fig. S3C).

### Structurally promiscuous clones dominate healthy human serum IgA1

By using SEC fractionation prior to IgA1 clonal profiling, we were able to extend the depth of our analysis considerably, whereby the resulting total serum IgA1 repertoires contained more than 1,400 unique clones per donor. For each of the nearly 3,000 unique endogenous IgA1 Fab clones identified in total from the two donors analyzed, we obtained individual SEC elution profiles and thus were able to determine each clone’s assembly distribution. All results from this analysis are summarized in SI Table S3.

The majority of unique clones were present solely as IgA1 monomers, with considerably fewer (but still >100 for each donor) found exclusively as J-coupled dimers (SI Table S3). In addition to these “extremes”, in both donors we also observed hundreds of unique clones that co-occurred in both mIgA and dIgA forms, readily evidenced by their appearance (green markers) at the same RPLC RT-Fab mass coordinates in the dIgA1 and mIgA1 repertoires shown in SI Fig. 3A and B. Thus, human serum does not simply contain two distinct IgA1 clonal repertoires corresponding to separate pools of monomers and J-coupled dimers. Instead, the “total” serum IgA1 repertoire is divided into three subpopulations of unique clones, hereafter referred to as “monomer-only”, “dimer-only”, and “co-occurring” and color-coded throughout this manuscript by yellow, blue, and green, respectively (Fig. 2C). We also define two assembly-specific “monomer” and “dimer” clonal repertoires (SI Fig. 3A and B, respectively). The relationship between these repertoires (total, monomer, dimer) and unique clone (monomer-only, dimer-only, co-occurring) classifications is illustrated schematically in Fig. 2C.

**Figure 3.**
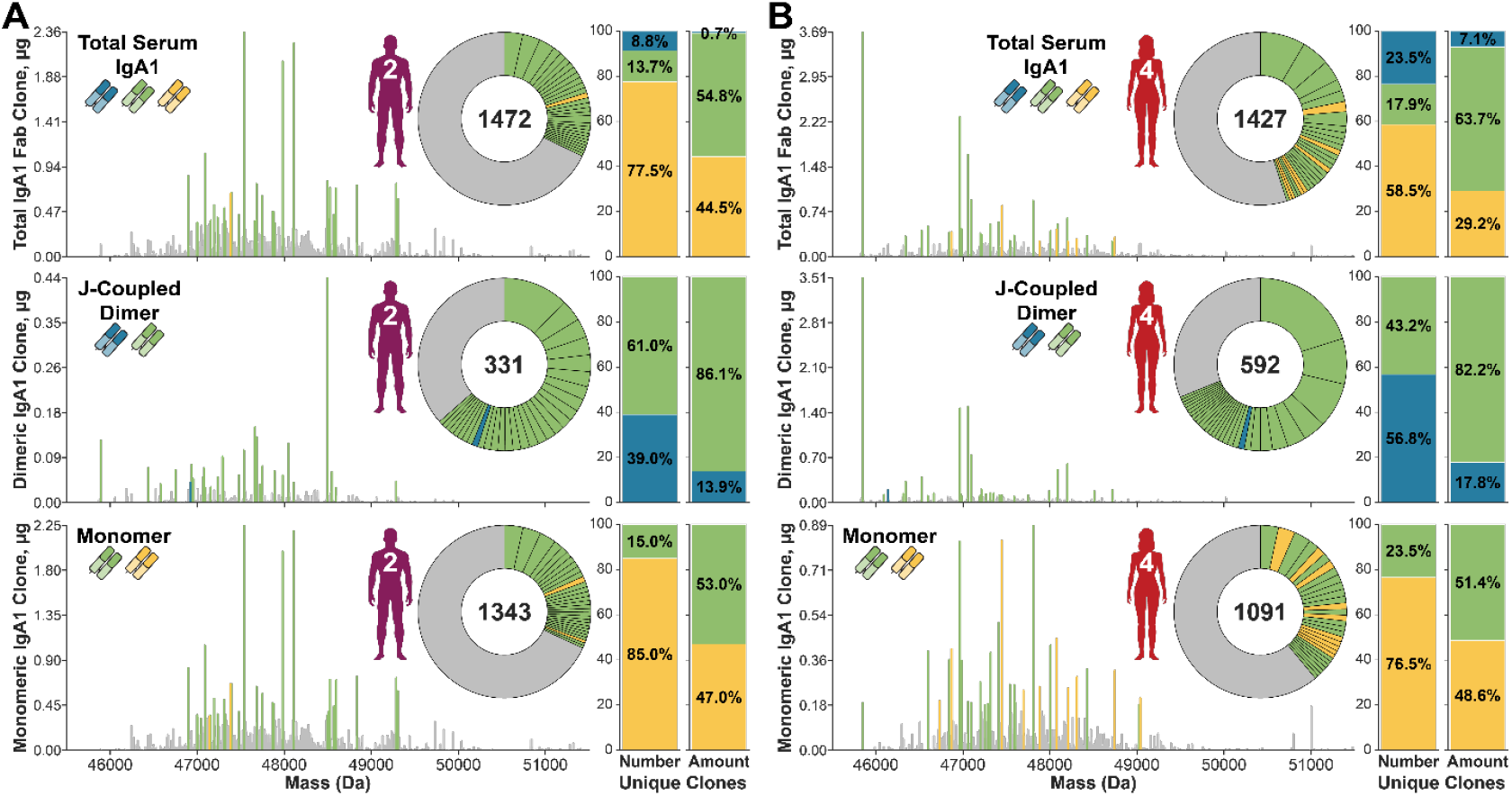
A small proportion of unique clones present in both monomer and J-coupled dimer forms dominates serum IgA1 repertoires. Total **(top)**, J-coupled dimer **(middle)**, and monomer **(bottom)** circulatory IgA1 clonal profiles for Donor 2 **(A)** and Donor 4 **(B)**. Each stick represents a unique IgA1 Fab clone, plotted according to its characteristic mass and abundance. The top 30 most abundant clones in each panel are annotated as monomer-only (yellow), dimer-only (blue) or co-occurring (green), with their fractional abundance indicated by the relative size of the colored wedge slices of the inset donut chart. All clones ranking outside the top 30 most abundant are represented by grey. The total number of unique IgA1 Fabs detected in each repertoire is indicated at the center of the inset. Bar charts to the right of each plot indicate the proportion of the total number of unique clones (**left** bar chart in each pair) and fractional abundance (**right** bar chart in each pair) of unique clones in the indicated circulatory IgA1 repertoire contexts.

Consistent with previous analyses of IgG1 and IgA1 Fab clonal profiles (***24-26***), both donors’ total and assembly-specific serum IgA1 repertoires are relatively simple, dominated by tens to hundreds of highly-abundant clones. In more detail, just ∼5% of clones collectively contribute ∼50% of the total IgA1 abundance in each repertoire (SI Fig. S4). Here, we highlight this trend holds true even though we detect significantly more distinct clones (∼3.5-fold, >1,400 per sample) with improved sensitivity owing to the additional SEC-based separation, as compared to previous IgA1 Fab clonal profiling in serum (***26***).

Remarkably, of the three types of unique clones revealed here, the most dominant clones are largely part of the co-occurring clonal population (Fig. 3). In the total serum IgA1 clonal repertoires of Donors 2 and 4 (Fig. 3A and B, respectively), by their numbers less than 20% of all unique clones are classified as co-occurring, and yet as a group their cumulative fractional abundance accounts for over 50% of the total circulatory IgA1.

Likewise, J-coupled dimer repertoires are dominated by clones co-occurring in the monomeric fraction rather than by those found exclusively as dIgA (Fig. 3). Although they contain fewer unique clones (SI Fig. S3C) and lower concentrations than the monomer and total serum IgA1 repertoires (SI Table S3), these J-coupled dimeric IgA1 repertoires are especially “simple”. Indeed, a staggering ∼60-70% of the total abundance is contributed by just the top 30 clones, i.e., fewer than one-tenth of all unique clones occurring in J-coupled dimer form (Fig. 3 and SI Fig. S4). This is particularly pronounced for Donor 4, for whom just the top seven most dominant clones account for half of the total J-coupled dimer amount (Fig. 3B). Despite Donor 4’s dimer-only clones (336) alone outnumbering Donor 2’s entire dimer repertoire (331), the composition of both donors’ top 30 dIgA clones is highly similar–i.e., all but one of these top dimer clones were also found as monomers without the J-chain. Thus Donor 4’s “extra” dimer-only clones have very low abundances (Fig. 3). Differences between the donors’ intact IgA assembly distributions observed by MP (Fig. 1B) therefore are not explained by the significant disparity in the number of unique clones.

### Assembly composition varies between individual IgA1 clones

Each donor’s cumulative IgA1 Fab abundance is differently distributed between the two assemblies (Fig. 2B) and among the three unique clone classes (Fig. 3). This indicates that the structurally-promiscuous IgA1s co-occur in different mIgA/dIgA assembly ratios. In particular, the J-coupled dimeric form must correspond to a much larger fraction of the total co-occurring clone abundance for Donor 4 than for Donor 2. Indeed, monomers account for >90% of the cumulative co-occurring clone abundance for Donor 2 (SI Fig. S5A). In contrast, in Donor 4 they are nearly equal with their J-coupled dimer counterparts overall (SI Fig. S5A). However, each unique co-occurring clone could have a varying distribution that does not match these overall trends, with some being predominantly monomeric, but other clones more dimeric (SI Fig. S5). We therefore next compared how the assembly ratios of each donor’s individual clones were distributed. For Donor 2, this distribution is heavily skewed toward the monomer-dominated end, compared to the relatively more even distribution for Donor 4 (SI Fig. S5B and Fig. 4A). Clearly, co-occurring clone assembly composition is not a donor-specific feature but instead appears to be clone-specific. This variation is best illustrated by plotting the SEC profiles of individual distinct IgA1 clones identified by their unique Fab mass-RT combination and quantified in each fraction (Fig. 4B). The appearance of a bimodal distribution is diagnostic for co-occurring clones in accordance with the established discrete elutions of intact J-coupled dimers and monomers. Examples selected from each donor’s top 100 most abundant IgA1 clones are shown in order of increasing J-coupled dimer fractional abundance (Fig. 4B). This reaches a maximum of ∼56% J-coupled dimer for Donor 2, while the top 100 most abundant IgA1 co-occurring clones from Donor 4 already illustrate the full extent of variation possible (Fig. 4B).

**Figure 4.**
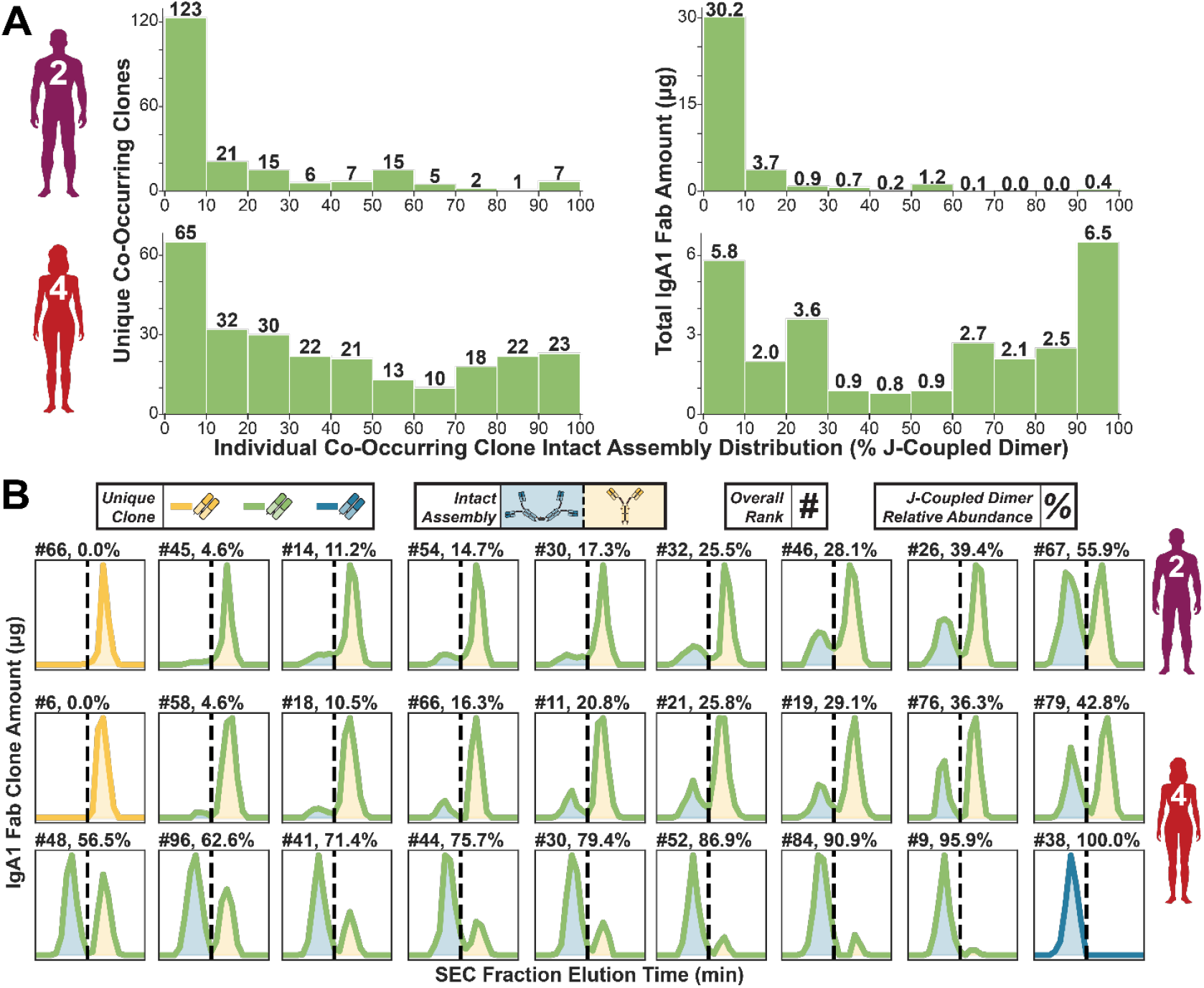
The assembly distribution of distinct co-occurring IgA1 clones varies widely. **(A)** Histograms depicting the assembly distribution of all unique co-occurring IgA1 clones in Donor 2 **(top)** and Donor 4 **(bottom)** according to number **(left)** and cumulative abundance **(right)**. The ten bins in each histogram are defined according to the relative intact assembly distribution (i.e., the fractional abundance of J-coupled dimer assemblies) of those clones. **(B)** SEC elution profiles of distinct co-occurring IgA1 Fab clones selected together with one monomer-only and one dimer-only clone (if applicable) from the top 100 overall most abundant clones in the total serum IgA1 repertoire of Donor 2 **(top row)** and Donor 4 **(bottom two rows)**. Each panel is annotated with the clone’s overall donor-specific ranking (#) and fractional J-coupled dimer abundance (%). Plots are shown in order of increasing J-coupled dimer fractional abundance (%). Line color indicates the corresponding unique clone type according to the color scheme in shown in the legend, with the area under the peaks shaded according to the discrete elution ranges of intact J-coupled dimers (blue) and monomers (yellow). Bimodal distributions directly indicate the clone’s co-occurrence in both forms.

We conclude from the data that each unique IgA1 clone in circulation could be present in a mixture of assembly states varying from solely monomeric to solely J-coupled dimeric (SI Fig. S5). In between these extremes and dominating the total serum repertoire are IgA1 clones that in circulation simultaneously occur both with and without J-chain, revealing that the structural promiscuity of human IgA is not just a bulk population property. The observation that each discrete IgA1 clone—defined by its unique IgA+ ASC origin— has a unique assembly distribution is, as far as we know, completely unprecedented and raises several new questions we address in the Discussion.

## Discussion

In the adaptive humoral immune response, binding of an antigen to the B-cell receptor induces naïve B cells to proliferate, undergo affinity maturation, and ultimately differentiate into an ASC capable of producing massive quantities of antigen-specific antibody. Each Ig sequence derived from a different B cell is not only unique but also directly reflects its clonal origins and unique antigen target (***32, 33***). In our protein-centric approach, each unique IgA1 clone is defined by its unique combination of Fab mass and RPLC RT signature, independent of the assembly origin(s). A view commonly presented in IgA literature is that J-coupled polymeric IgA and monomeric (i.e., non-J-coupled) IgA—which dominate mucosal and systemic sites, respectively—originate from two distinct populations of IgA+ ASCs in which J-chain is expressed (J+) or absent (J−) (***30***). Accordingly, monomers and J-coupled dimers of IgA should correspond to two distinct clonal repertoires with no overlap in cellular origins or antigen binding, which clearly does not align with the current findings. Moreover, foundational early experiments supporting the persistent view that circulatory IgA-producing ASCs in the bone marrow are J− (***34***) likely suffered from technical limitations and inadvertently under-estimated the amount of J-chain, as summarized nicely by Castro and Flajnik (***30***). Based on more recent advances, the J-chain is nowadays more commonly considered to be a marker of B cell differentiation into ASCs (***35-39***), although there are still many open questions regarding how J-chain expression and incorporation into IgA is regulated (***30***). However, *in vitro* experiments have reliably shown that IgA can be produced recombinantly as an array of mIgA, dIgA, and pIgA assemblies which varies depending on the relative amounts of intracellular J-chain, IgA heavy chain, and light chain (***40, 41***).

These contradicting views illustrate that there are still many open questions around IgA (***30***), and our findings here suggest the human IgA system is much more complex than has been long considered. We hereby propose a model in which there are three populations of IgA+ ASCs that contribute to the serum IgA pool in healthy individuals (Fig. 5A). The relative amount of J-chain to IgA expression in the ASC dictates the assembly distribution produced according to three regimes delimited by a minimum threshold for formation of dIgA and an upper limit for mIgA production. At the extremes, J^low^ and J^high^ IgA+ ASCs exclusively produce mIgA and dIgA, respectively. Here, we provide novel evidence supporting the existence of J^mid^ IgA+ ASCs that produce IgA1 clones in varied monomer/J-coupled dimer ratios (Fig. 5A). The high prevalence of IgA1 clones produced in both mIgA1 and dIgA1 forms—in both donors, regardless of the large differences between their intact assembly distributions—suggests the same population of IgA+ ASCs can indeed be engaged in the simultaneous synthesis of both J-devoid monomeric and J-coupled dimeric IgA (***42, 43***).

**Figure 5.**
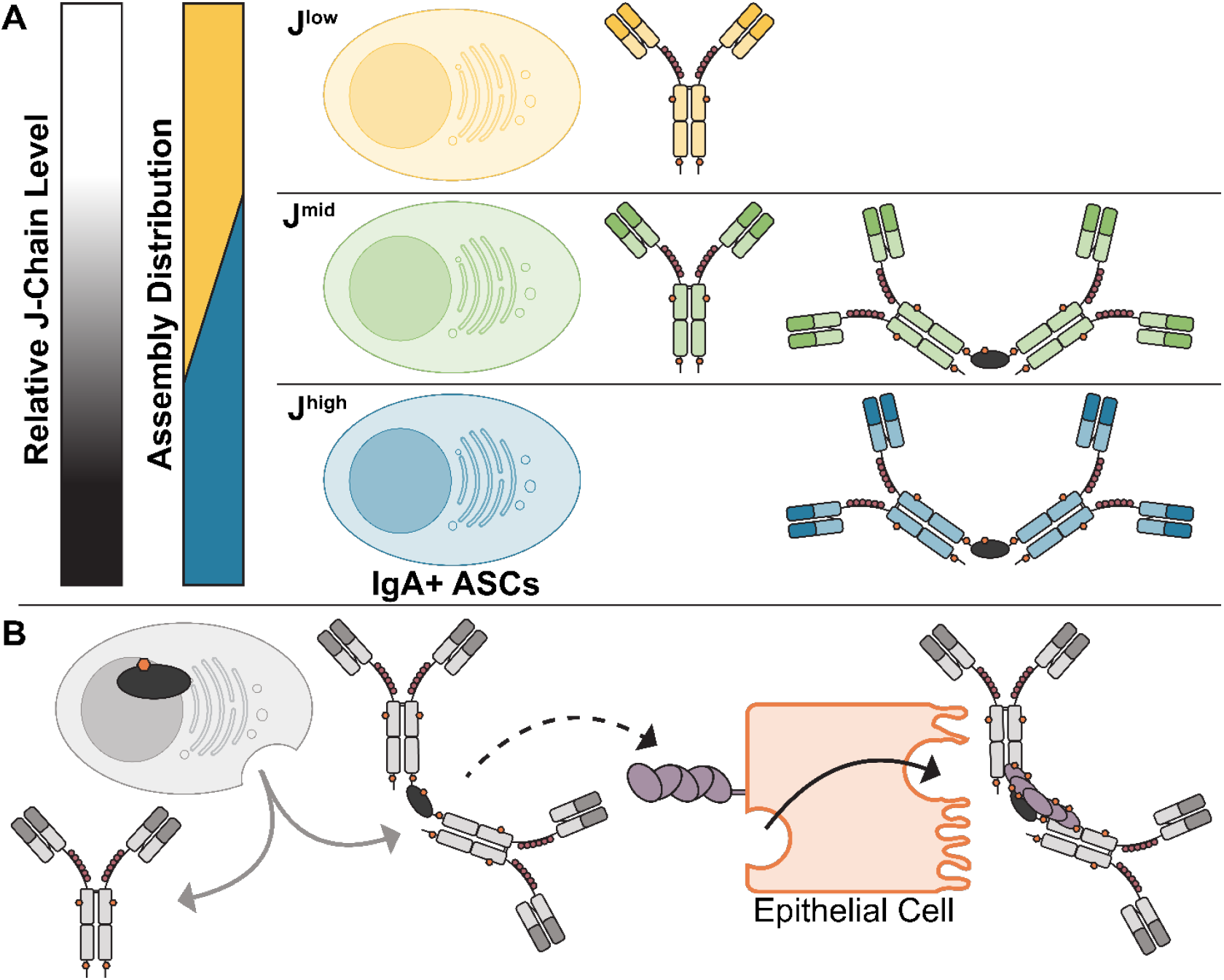
Graphical summary. **(A)** IgA+ ASCs co-express variable amounts of J-chain, determining the fractional abundance of J-coupled dIgA. Three distinct regimes are proposed in the context of the two assembly states and three clonal populations of human circulatory IgA1 in healthy individuals observed by assembly-specific IgA1 Fab clonal profiling. **(B)** Structural promiscuity with the J-chain can also result in shared clonality between circulatory and mucosal sites. Monomers lack the J-chain necessary for pIgR-mediated transport across the mucosal epithelium and thus enter circulation. A variable proportion of J-coupled dimer produced by the same IgA+ ASC may also enter circulation, or, if produced locally at a mucosal site, may be transported into the mucosa as SIgA.

### Implications of structural promiscuity on human IgA function

Each molecule of a unique IgA clone recognizes the same antigen determined by the Fab variable region, irrespective of the antibody being in the mIgA or dIgA form. These two assembly states of the same unique clone may, however, elicit very different immunological effects for the same target (***18, 44, 45***). For example, circulating mIgA immune complexes in particular are noted for inducing an immediate pro-inflammatory response by cross-linking FcαRI (CD89) (***19, 46, 47***). In contrast, in mucosal immune contexts, SIgA instead is known to be anti-inflammatory and contribute to homeostatic conditions by utilizing anti-inflammatory mechanisms such as immune exclusion and neutralization of pathogens. Here, dIgA is even able to intercept and neutralize invading viruses intracellularly during pIgR-mediated transepithelial transport to the mucosal lumen, which is inaccessible to mIgA due to the absence of J-chain (***47-50***). The distinct structures and functions of the different molecular forms of human IgA also support the conclusion from our data that structural promiscuity varies for each unique IgA1 clone (Fig. 4 and SI Fig. S5). Moreover, for each clone the ratio of mIgA1 to dIgA1 may be variable over time, potentially reflecting differences in immune response timing. Generation of high-affinity antigen-specific antibodies requires weeks, restricting initial responses to low-affinity antibodies. In these low-affinity responses, rapid production of exclusively J-coupled dIgA could compensate due to its enhanced avidity and neutralization capacity (***50-52***). Interestingly, antigen-specific circulatory IgAs produced early in response to viral or bacterial infection have indeed been described as predominantly polymeric and typically later replaced by a persistent monomer response (***10, 17***).

Notwithstanding that many factors may contribute to the observed assembly distribution for each clone (***4, 30, 53***), the data shown here point toward a significant role for the J-chain-containing immunocomplexes in the circulatory IgA response. From a biotherapeutic development perspective, our findings could have significant impact. Although the field of antibody therapeutics is heavily dominated by recombinant antibodies designed by using an IgG (mostly IgG1) template, there is a growing interest to develop the class of IgA as potential alternative biotherapeutic (***54***). With not only multiple N-glycans but also O-glycans present, IgA is one of the most heavily glycosylated human antibodies and structurally more complicated than IgG. However, the fact that IgA could be made into distinct modalities—i.e., mIgA and dIgA, with known differences in stability, clearance rates, and especially the higher avidity and enhanced neutralization capacity reported for J-coupled dimers—makes IgA-based therapeutics an exciting venue to explore (***50, 55***).

### Implications for shared clonality between serum and mucosal IgA

As shown here and in our earlier studies involving more donors (***24-26, 56***), clonal repertoires are entirely unique when comparing serum (or milk) samples from different donors (i.e., nearly every IgG1 or IgA1 Fab detected in each donor has its own unique mass/RT signature). However, we previously observed a high level of shared clonality between IgA1 from serum and breastmilk of the same donor (***26***), initially presumed to correspond solely to clones present in circulation as J-coupled dimers. From the structural promiscuity we have uncovered here, we now know the same distinct IgA1 found as SIgA at mucosal sites may not be exclusively dimeric in serum (Fig. 5B). Adapting our approach to enable the comparison of assembly-specific IgA clonal repertoires in serum with those found in mucosal compartments is thus of high interest, as it should provide a clearer picture of the cellular origins, trafficking, clonality, and functions of the human IgA system (***57***). Blood is the crossroad of not just circulatory Ig but also migrating plasmablasts and plasma cells (PCs), which may be actively secreting antibodies (***58, 59***). Notably, of these circulating ASCs, 80% are reportedly IgA+ (***60***). Several reports have also provided evidence suggesting stable, significant contributions to serum IgA by cells originally induced at mucosal sites (***39, 60-62***).

Together this suggests each donor’s circulatory IgA clonal repertoire contains a broad, diverse mixture of antigen specificities, from all different inductive sites (Fig. 5B). We propose that J-chain structural promiscuity enables the structural and functional diversification of human IgA, exemplified by co-occurring clones. Too much clonal diversity risks autoimmunity, with negative health consequences especially in gut mucosal contexts home to both the largest population of IgA+ PCs and vast communities of microbes. By essentially copying the same IgA1 but changing the structural format, antibody diversity is increased while maintaining a restricted sequence repertoire. Considering the extensive, highly abundant distribution of human IgA throughout the body in many different compartments where diverse functions and signaling outcomes are required, this could thus, for example, enable serum IgA to function as a failsafe against sepsis if the mucosal epithelial barrier is breached (***19, 63, 64***).

### Outlook

Taken together, our findings paint a broader, more complicated picture of human serum IgA (Fig. 5) in complete contrast with what is generally depicted in the literature (where the contributions of circulatory J-coupled IgA dimers are often ignored). This could result in overlooking nearly one-third of serum IgA, including many highly-abundant unique clones, and potentially arriving at flawed or altogether-erroneous conclusions about the characteristics of human circulatory IgA if molecular and/or clonal composition is not directly assessed. The ability of the same distinct IgA1 antibody to occur both with and without the J-chain necessarily increases the complexity of serum IgA. This structural promiscuity of IgA with the J-chain introduces challenges, but more importantly it also inspires many exciting new questions to address in the future in order to fully tease apart the intriguing system of human IgA.

## Materials and Methods

All donor samples were obtained with approval of the ethical advisory board and with written informed consent of the donors where applicable, as described in the SI Appendix. A detailed description of all materials and methods used, including affinity purification, offline size exclusion chromatography, and generation of IgA1 Fabs, as well as for all mass photometry and mass spectrometry-based analyses, is provided in the Supplementary Materials and Methods section of the SI Appendix.

## Data, Materials, and Software Availability

All study data are included in the article and/or SI Appendix.

## Acknowledgments

A.D.R. acknowledges support from the European Molecular Biology Organization through a Long-Term Postdoctoral Fellowship (ALTF 371-2022). A.J.R.H. acknowledges support from the Netherlands Organization for Scientific Research (NWO) through the Spinoza Award (SPI.2017.028). We thank Genmab (Utrecht, NL) for their long-term financial support of our work and for the gift of recombinant monomeric anti-CD20 and anti-MET IgA1 mAbs used as internal standards in this study. We also acknowledge Sanquin Research (Amsterdam, NL) for providing serum samples.

## References

1. J. M. Woof, J. Mestecky, “Mucosal Immunoglobulins” in Mucosal Immunology (Fourth Edition), J. Mestecky, W. Strober, M. W. Russell, B. L. Kelsall, H. Cheroutre, B. N. Lambrecht, Eds. (Academic Press, 2015), pp. 287–324.

2. P. de Sousa-Pereira, J. M. Woof, IgA: Structure, Function, and Developability. Antibodies 8, 57 (2019).

3. M. E. Conley, D. L. Delacroix, Intravascular and Mucosal Immunoglobulin A: Two Separate but Related Systems of Immune Defense? Ann. Intern. Med. 106, 892–899 (1987).

4. M. L. Matsumoto, Molecular Mechanisms of Multimeric Assembly of IgM and IgA. Annu. Rev. Immunol. 40, 221–247 (2022).

5. M. E. Koshland, The Coming of Age of the Immunoglobulin J Chain. Annu. Rev. Immunol. 3, 425–453 (1985).

6. V. Sørensen, I. B. Rasmussen, V. Sundvold, T. E. Michaelsen, I. Sandlie, Structural requirements for incorporation of J chain into human IgM and IgA. Int. Immunol. 12, 19–27 (2000).

7. N. Oskam et al., CD5L is a canonical component of circulatory IgM. Proc. Natl. Acad. Sci. U.S.A. 120, e2311265120 (2023).

8. N. Kumar, C. P. Arthur, C. Ciferri, M. L. Matsumoto, Structure of the secretory immunoglobulin A core. Science 367, 1008–1014 (2020).

9. Y. Wang, J. Xiao, Recent advances in the molecular understanding of immunoglobulin A. FEBS J. 291, 3597–3603 (2024).

10. M. W. Russell, C. Lue, A. W. L. van den Wall Bake, Z. Moldoveanu, J. Mestecky, Molecular heterogeneity of human IgA antibodies during an immune response. Clin. Exp. Immunol. 87, 1–6 (2008).

11. V. Snoeck, I. R. Peters, E. Cox, The IgA system: a comparison of structure and function in different species. Vet. Res. 37, 455–467 (2006).

12. U. Steffen et al., IgA subclasses have different effector functions associated with distinct glycosylation profiles. Nat. Commun. 11, 120 (2020).

13. S. Pan et al., Each N-glycan on human IgA and J-chain uniquely affects oligomericity and stability. Biochim. Biophys. Acta Gen. Subj. 1868, 130536 (2024).

14. A. Rifai, K. Fadden, S. L. Morrison, K. R. Chintalacharuvu, The N-Glycans Determine the Differential Blood Clearance and Hepatic Uptake of Human Immunoglobulin (Ig)A1 and IgA2 Isotypes. J. Exp. Med. 191, 2171–2182 (2000).

15. D. Falck et al., IgA displays site- and subclass-specific glycoform differences despite equal glycoenzyme expression. Cell Commun. Signal. 23, 92 (2025).

16. S. Jackson, Z. Moldoveanu, J. Mestecky, “Collection and Processing of Human Mucosal Secretions” in Mucosal Immunology (Fourth Edition), J. Mestecky, W. Strober, M. W. Russell, B. L. Kelsall, H. Cheroutre, B. N. Lambrecht, Eds. (Academic Press, 2015), pp. 2345–2353.

17. O. Pabst, E. Slack, IgA and the intestinal microbiota: the importance of being specific. Mucosal Immunol. 13, 12–21 (2020).

18. S. K. Davis, K. J. Selva, S. J. Kent, A. W. Chung, Serum IgA Fc effector functions in infectious disease and cancer. Immunol. Cell Biol. 98, 276–286 (2020).

19. K. W. Leong, J. L. Ding, The Unexplored Roles of Human Serum IgA. DNA Cell Biol. 33, 823–829 (2014).

20. J. Radl, A. C. W. Swart, J. Mestecky, The Nature of the Polymeric Serum IgA In Man. Proc. Soc. Exp. Biol. Med. 150, 482–484 (1975).

21. D. L. Delacroix et al., Changes in size, subclass, and metabolic properties of serum immunoglobulin A in liver diseases and in other diseases with high serum immunoglobulin A. J. Clin. Invest. 71, 358–367 (1983).

22. A. R. Rees, Understanding the human antibody repertoire. mAbs 12, 1729683 (2020).

23. S. L. Nutt, P. D. Hodgkin, D. M. Tarlinton, L. M. Corcoran, The generation of antibody-secreting plasma cells. Nat. Rev. Immunol. 15, 160–171 (2015).

24. A. Bondt, K. A. Dingess, M. Hoek, D. M. H. van Rijswijck, A. J. R. Heck, A Direct MS-Based Approach to Profile Human Milk Secretory Immunoglobulin A (IgA1) Reveals Donor-Specific Clonal Repertoires With High Longitudinal Stability. Front. Immunol. 12, 789748 (2021).

25. A. Bondt et al., Human plasma IgG1 repertoires are simple, unique, and dynamic. Cell Syst. 12, 1131-1143.e1135 (2021).

26. K. A. Dingess et al., Identification of common and distinct origins of human serum and breastmilk IgA1 by mass spectrometry-based clonal profiling. Cell. Mol. Immunol. 20, 26–37 (2023).

27. W. H. Kutteh, S. J. Prince, J. Mestecky, Tissue origins of human polymeric and monomeric IgA. J. Immunol. 128, 990–995 (1982).

28. R. Iversen et al., Strong Clonal Relatedness between Serum and Gut IgA despite Different Plasma Cell Origins. Cell Rep. 20, 2357–2367 (2017).

29. V. Demichev et al., A time-resolved proteomic and prognostic map of COVID-19. Cell Syst. 12, 780-794.e787 (2021).

30. C. D. Castro, M. F. Flajnik, Putting J Chain Back on the Map: How Might Its Expression Define Plasma Cell Development? J. Immunol. 193, 3248–3255 (2014).

31. A. D. Rolland, A. J. R. Heck, “Analytical Challenges in Analyzing IgM Biotherapeutics” in Characterizing Biotherapeutics: Analytical Methods for Diverse Modalities, J. R. Lill, W. Sandoval, Eds. (Wiley, 2025), pp. 29–54.

32. H. Li et al., Mucosal or systemic microbiota exposures shape the B cell repertoire. Nature 584, 274–278 (2020).

33. J. Tellier et al., Unraveling the diversity and functions of tissue-resident plasma cells. Nat. Immunol. 25, 330–342 (2024).

34. J. Mestecky, J. R. McGhee, Immunoglobulin A (IgA): Molecular and Cellular Interactions Involved in IgA Biosynthesis and Immune Response. Adv. Immunol. 40, 153–245 (1987).

35. A. Q. Xu, R. R. Barbosa, D. P. Calado, Genetic timestamping of plasma cells in vivo reveals tissue-specific homeostatic population turnover. eLife 9, e59850 (2020).

36. D. Morgan, V. Tergaonkar, Unraveling B cell trajectories at single cell resolution. Trends Immunol. 43, 210–229 (2022).

37. M. Duan et al., Understanding heterogeneity of human bone marrow plasma cell maturation and survival pathways by single-cell analyses. Cell Rep. 42, 112682 (2023).

38. N. J. M. Verstegen et al., Single-cell analysis reveals dynamics of human B cell differentiation and identifies novel B and antibody-secreting cell intermediates. eLife 12, e83578 (2023).

39. F. Vaz et al., Distinct systemic and gut IgA responses to bacteria of the human upper gastrointestinal tract. bioRxiv [Preprint] (2025). 10.1101/2025.07.01.662496 (accessed 21 October 2025).

40. R. M. E. Parkhouse, E. D. Corte, Biosynthesis of immunoglobulin A (IgA) and immunoglobulin M (IgM). Control of polymerization by J chain. Biochem. J. 136, 607–609 (1973).

41. T. N. Lombana et al., Production, characterization, and in vivo half-life extension of polymeric IgA molecules in mice. mAbs 11, 1122–1138 (2019).

42. A. Tarkowski et al., Cellular origins of human polymeric and monomeric IgA: enumeration of single cells secreting polymeric IgA1 and IgA2 in peripheral blood, bone marrow, spleen, gingiva and synovial tissue. Clin. Exp. Immunol. 85, 341–348 (1991).

43. Z. Moldoveanu, M. L. Egan, J. Mestecky, Cellular origins of human polymeric and monomeric IgA: intracellular and secreted forms of IgA. J. Immunol. 133, 3156–3162 (1984).

44. P. J. Gleeson, N. O. S. Camara, P. Launay, A. Lehuen, R. C. Monteiro, Immunoglobulin A Antibodies: From Protection to Harmful Roles. Immunol. Rev. 328, 171–191 (2024).

45. I. S. Hansen, D. L. P. Baeten, J. den Dunnen, The inflammatory function of human IgA. Cell. Mol. Life Sci. 76, 1041–1055 (2019).

46. G. Vidarsson et al., Activity of Human IgG and IgA Subclasses in Immune Defense Against Neisseria meningitidis Serogroup B1. J. Immunol. 166, 6250–6256 (2001).

47. M. M. J. van Gool, M. van Egmond, IgA and FcαRI: Versatile Players in Homeostasis, Infection, and Autoimmunity. Immunotargets Ther. 9, 351–372 (2021).

48. D. Sinha, M. Yaugel-Novoa, L. Waeckel, S. Paul, S. Longet, Unmasking the potential of secretory IgA and its pivotal role in protection from respiratory viruses. Antiviral Res. 223, 105823 (2024).

49. D. Sterlin et al., IgA dominates the early neutralizing antibody response to SARS-CoV-2. Sci. Transl. Med. 13, eabd2223 (2021).

50. Z. Wang et al., Enhanced SARS-CoV-2 neutralization by dimeric IgA. Sci. Transl. Med. 13, eabf1555 (2021).

51. M. A. Maurer et al., Glycosylation of Human IgA Directly Inhibits Influenza A and Other Sialic-Acid-Binding Viruses. Cell Rep. 23, 90–99 (2018).

52. S. Saito et al., IgA tetramerization improves target breadth but not peak potency of functionality of anti-influenza virus broadly neutralizing antibody. PLOS Pathog. 15, e1007427 (2019).

53. E. Xiong et al., MZB1 promotes the secretion of J-chain–containing dimeric IgA and is critical for the suppression of gut inflammation. Proc. Natl. Acad. Sci. U.S.A. 116, 13480–13489 (2019).

54. D. Sterlin, G. Gorochov, When Therapeutic IgA Antibodies Might Come of Age. Pharmacology 106, 9–19 (2020).

55. K. Göritzer et al., Recombinant neutralizing secretory IgA antibodies for preventing mucosal acquisition and transmission of SARS-CoV-2. Mol. Ther. 32, 689–703 (2024).

56. A. Bondt et al., Into the Dark Serum Proteome: Personalized Features of IgG1 and IgA1 Repertoires in Severe COVID-19 Patients. Mol. Cell. Proteomics 23, 100690 (2024).

57. W. Meng et al., An atlas of B-cell clonal distribution in the human body. Nat. Biotechnol. 35, 879–884 (2017).

58. M. Perez-Andres et al., Human peripheral blood B-cell compartments: A crossroad in B-cell traffic. Cytometry B Clin. Cytom. 78B, S47–S60 (2010).

59. B. Isho, A. Florescu, A. A. Wang, J. L. Gommerman, Fantastic IgA plasma cells and where to find them. Immunol. Rev. 303, 119–137 (2021).

60. H. E. Mei et al., Blood-borne human plasma cells in steady state are derived from mucosal immune responses. Blood 113, 2461–2469 (2009).

61. S. J. Keppler, M. C. Goess, J. M. Heinze, The Wanderings of Gut-Derived IgA Plasma Cells: Impact on Systemic Immune Responses. Front. Immunol. 12, (2021).

62. A. Lemke et al., Long-lived plasma cells are generated in mucosal immune responses and contribute to the bone marrow plasma cell pool in mice. Mucosal Immunol. 9, 83–97 (2016).

63. J. R. Wilmore et al., Commensal Microbes Induce Serum IgA Responses that Protect against Polymicrobial Sepsis. Cell Host Microbe 23, 302-311.e303 (2018).

64. J. Schlechte, I. Skalosky, M. B. Geuking, B. McDonald, Long-distance relationships - regulation of systemic host defense against infections by the gut microbiota. Mucosal Immunol. 15, 809–818 (2022).

