## Supporting Information for "Structural promiscuity in the human circulatory IgA1 clonal repertoire"

**This PDF file includes:**

Supplementary Materials and Methods
Figures S1 to S5
Tables S1 to S3
SI References

### Supplemental Materials and Methods

#### Healthy donor serum/plasma and IgA1 mAb materials

*Donor samples.* Individual serum samples from anonymous healthy volunteers (Donors 1, 3, and 4) were collected after obtaining an informed consent, approved by the ethical approval was obtained from the Sanquin Ethical Advisory Board before participation. The individual $K_2$EDTA plasma sample (Donor 2) was obtained commercially from Zen-Bio (Durham, North Carolina, USA) and was collected from a healthy volunteer undergoing an elective procedure in compliance with ethical regulations and with informed consent. All samples were stored at −80 °C until further analysis. Donor demographic information can be found in Table S1.

*IgA1 monoclonal antibodies (mAbs).* Two recombinant monomeric IgA1 mAbs, 7D8-IgA1 (anti-CD20) and 5D5v2-IgA1 (anti-MET), were used as internal standards to allow intensity-based quantitation of endogenous IgA1 Fab clones. These were provided as a gift by Genmab (Utrecht, NL).

#### Affinity purification of intact circulatory IgA assemblies from individual donor sera and plasma

*Preparation of affinity resin.* Affinity purification of all intact IgA assemblies from whole donor serum or plasma directly was achieved using CaptureSelect IgA Affinity Matrix (Thermo Scientific) which binds all subclasses/allotypes and all molecular forms of human IgA (including dimeric and secretory) via the Fc domain. For each sample, an appropriate volume of affinity resin slurry (1:1 beads:buffer) was pipetted into a Pierce screw cap spin column (Thermo Scientific) and then washed repeatedly by adding 150-200 µL PBS followed by centrifugation for 1 min at 500 × *g*. Following at least three wash cycles, the columns were plugged and the sample was added.

*Isolation of IgA from plasma/serum.* For affinity purification of intact IgA assemblies directly from whole donor plasma/serum for subsequent analysis by mass photometry and/or size exclusion chromatography, 20 µL plasma/serum was added together with 150 µL PBS. The samples were then incubated on an Eppendorf thermal shaker for 1 hr at room temperature and 750 rpm to allow binding of all IgA to the affinity resin beads. Following this incubation period, the plugs were removed from the spin columns, and the flow-through was collected by centrifugation (1 min, 500 × *g*).

*Elution of intact IgA assemblies.* For purification of intact IgA assemblies for subsequent analysis by mass photometry and/or size exclusion chromatography, the beads were then washed twice with PBS followed by two more wash steps with Milli-Q water to remove all unbound proteins. The affinity-captured IgA was then eluted from the beads in the plugged spin columns by adding 100 µL 0.4% formic acid (HCOOH) in Milli-Q water and incubating 10 min at room temperature while shaking at 750 rpm. Then the plug was removed and the eluate containing the intact IgA assemblies was collected as the flow-through into LoBind tubes following centrifugation (1 min, 1000 × *g*). Eluted samples were neutralized by adding 20 µL Tris base (1 M, pH 8.0) and stored briefly at 4 °C until further analysis of the intact IgA assemblies by single-molecule mass photometry or size exclusion chromatography.

#### Mass photometry of affinity-purified intact IgA assemblies

*Preparation of coverslips.* Borosilicate glass microscope coverslips (24 × 50 mm) were cleaned by sonication in isopropanol followed by Milli-Q water, and this process was repeated for a total of four rounds. A nitrogen gas spray gun was used to dry the clean coverslips and remove any dust particles. Silicone cell culture gaskets were cut into sets of six wells (1 mm deep × 3 mm in diameter), rinsed with isopropanol followed by Milli-Q water, dried with a Kimwipe, and then placed in the center of a clean, dry coverslip. Prepared coverslips were stored in a closed container to prevent dust accumulation.

*Data acquisition.* All mass photometry (MP) experiments were performed on a Refeyn OneMP mass photometer (Refeyn Ltd., Oxford, UK). Prior to each measurement, 12 µL of PBS was placed in the unused coverslip well to re-focus the instrument on the glass-liquid interface, after which 3 µL of pre-diluted sample was added and mixed in the droplet via pipetting. Intact IgA affinity-purified from each of the four healthy donor plasma/serum samples was diluted with PBS accordingly to ensure a final concentration of ~10-100 nM in the droplet, with pre-diluted samples stored on ice for several minutes between the initial and subsequent 1:5 in-droplet dilutions. Calibration was performed using a mixture prepared in-house in PBS consisting of four proteins with accurate masses previously determined by mass spectrometry: IgG half-body (IgG4Δhinge-L368A), 73 kDa; IgG1-Campath, 149 kDa; apoferritin, 483 kDa; and GroEL, 800 kDa. All measurements were recorded in AcquireMP software (Refeyn) for 60 s in the medium field-of-view.

_Data processing and analysis._ Data were processed with DiscoverMP (Refeyn) to convert the measured ratiometric contrast measured in association with each "event" (i.e., a single molecule detected upon landing on the glass coverslip (***1***)) to mass based on the calibrant values and then exported for further analysis and plotting using an in-house python library. Mass histogram peaks were fit with Gaussian distributions to determine the mass of each species and the total number of detected molecules, enabling quantitative comparison between the assigned monomeric and J-coupled dimeric IgA assembly populations. These unadjusted values were used to calculate the fraction of total intact IgA assemblies and ratio of monomers to J-coupled dimers listed in Table S2. Considering each J-coupled dimer contains two IgA subunits, or twice as much IgA heavy chain, as a monomer, we doubled the number of detected J-coupled dimer assemblies without scaling the monomer assembly population. These adjusted values were used to determine the fraction of total serum IgA present as monomers and J-coupled dimers for each donor sample shown in Figure 1B of the main text and also listed in Table S2.

### Bottom-up proteomics sample preparation

All samples analyzed by mass spectrometry-based bottom-up proteomics (both unfractionated and SEC-fractionated single-donor plasma/serum and affinity-purified intact IgA samples) were prepared in the same way. Each donor plasma/serum sample was first diluted as necessary to achieve a total protein content of 5-7 µg in a minimal volume of 2 µL, to which 78 µL of lysis buffer was added. The lysis buffer contained: 1%(w/v) sodium deoxycholate (SDC), 10 mM tris(2-carboxyethyl)phosphine hydrochloride (TCEP), 100 mM Tris-HCl, 40 mM chloroacetamide (CAA), pH 8.5. Samples were then subjected to denaturation and alkylation at 95 °C for 5 min, followed by 2-fold dilution with Tris-HCl buffer. Trypsin (Sigma-Aldrich) and Lys-C (Wako) enzymes were added to each sample at enzyme-to-protein ratios of 1:50 and 1:75, respectively, for overnight digestion at 37 °C. The following day, trifluoroacetic acid (TFA) was added to each sample accordingly to reach a final concentration of roughly 1-4%(v/v) formic acid to stop the enzymatic digest and precipitate the SDC. Acidified samples were then centrifuged for 20 min at 20,000 × *g* to pellet the precipitate. The supernatant containing the peptides was collected and further diluted 40-fold with 1% Tris-HCl. All samples were stored at −20 °C if not immediately subjected to LC-MS/MS analysis.

### Proteomics analysis of affinity-purified IgA and total serum/plasma of individual healthy donors

_LC-MS/MS._ Peptide samples (20 µL) were loaded onto Evosep Pure tips according to Evosep manufacturer protocol. Peptides were separated using the 60 SPD method on an Evosep One LC system (Evosep, Odense, Denmark) with an EV-1109 analytical column (ReproSil Saphir C18, 1.5 µm beads by Dr Maisch; 8 cm length × 150 µm i.d.; Evosep). Gradient elution was achieved using mobile phases A (0.1% HCOOH in Milli-Q water) and B (0.1% HCOOH in $CH_3CN$). The LC was directly coupled to a Bruker timsTOF HT trapped ion mobility separation (TIMS) quadrupole time-of-flight (TOF) mass spectrometer (Bruker Daltonics, Bremen, Germany), and MS analysis was performed using dia-PASEF which combines parallel accumulation-serial fragmentation (PASEF) technology with data-independent acquisition (DIA) as previously described (***2***). MS1 scans were acquired in the range of 300-1,200 m/z with 12 dia-PASEF scans and 8.3% duty cycle. The ion mobility range was set from 1/K0 = 1.6 to 0.6 Vs/cm$^2$, and the ion accumulation and ramp times were set to 100 ms.

_Data processing and statistical analysis._ Raw files were searched using DIA-NN (version 1.8.1) in library-free mode (***3***). Trypsin was selected as the protease and two missed cleavages were tolerated. Cysteine carbamidomethylation and methionine oxidation were allowed. The mass accuracy was set to 20 and the MS1 accuracy to 10. The unrelated runs, isotopologues, match-between-runs, and no shared spectra options were enabled, whereas the heuristic inference was disabled. The protein interface was set to Genes and the IDs, RT and IM profiling was used for the library generation. The FDR was kept at the default 1%. The reviewed UniProt human protein database (canonical and isoforms) was used, with ~42,000 entries (Release number 2023_09) together with a manually curated contaminants database of around 150 entries. The main report of DIA-NN was filtered from the contaminants and from proteins with less than 2 identified peptides and used for further analysis. The abundance of each protein was calculated based on MaxLFQ values using the diann_maxlfq function of the DIAgui package (https://github.com/mgerault/DIAgui) (***4***). The Q.Value, Lib.Q.Value, and Lib.PG.Q.Value were each set to 1%. Lastly, proteins describing the variable domain of immunoglobulins were omitted.

*Determination of subclass and assembly distribution.* After processing and filtering, label-free quantitation (LFQ) values (Genes.MaxLFQ) were used to determine relative abundances of all proteins of interest. The IgA subclass distribution was determined from the unique peptides only (IGHA1 and IGHA2), depicted in Fig. S1B.

We also analyzed a publicly available plasma proteomics dataset (***5***) of 687 sampling points in aggregate from 139 patients, measured using data-independent acquisition mass spectrometry methods and analyzed with DIA-NN. Thus these data were obtained using similar approaches we employed in all MS-based proteomics analyses in this study, enabling comparison between results. Because the fractional abundance of J-coupled dimer assemblies determined from MP measurement was elevated in Donor 4 compared to literature, and considerable variation was found among our small cohort, we wanted to further investigate the distribution of circulatory IgA assemblies present in other individual human donors. In particular, we wanted to better benchmark the compositional landscape of healthy human IgA in serum to assess whether Donor 4 may or may not represent the upper limit of J-coupled dimer abundance and determine the "normal" (average) assembly composition we should expect.

We analyzed the protein quantities included in Table S8 of the Supplemental Information of this article (***5***) reported for IgA (IGHA1, IGHA2, IGHA1;IGHA2) in addition to JCHAIN, IGHM, and CD5L. All J-chain must be in complex with IgM or IgA. IgM occurs exclusively as pentamers (10 heavy chains) containing 1 J-chain and 1 CD5L. IgA is predominantly monomeric in circulation, but J-coupled dimers are also present. Thus we assigned the total J-chain detected abundance with priority to IgM, with the remainder representing the fraction of J-chain that must be associated with IgA. This in turn allows calculation of the fractional abundance of IgA present as J-coupled dimers and monomers lacking J-chain.

Due to uncertainty regarding the IgA2 allotype sequences used in the search (which will affect how the values are distributed between the two unique and one shared gene group) and considering that IgA1 dominates circulation anyway, we determined total IgA1 (IGHA1 + IGHA1;IGHA2) and used that instead of all IgA. The fraction of J-chain associated with IgM was calculated two ways (IGHM/10 and CD5L/1), with each value then subtracted from JCHAIN. The two remainders, representing the fraction of J-chain associated with IgA, were multiplied by 4 to determine the fraction of IGHA1 present in J-coupled dimers. These calculated values are plotted in Fig. S1C, with those corresponding to the publicly available plasma proteomics dataset (***5***) (N = 687 sampling points from 139 individuals) depicted in the histogram. For each of the four individuals included in the present study, both calculated values (arising from the estimation of IgM-associated J-chain as IGHM/10 and separately as CD5L/1) are shown as triangles with the color corresponding to that shown for Donors 1-4 in Fig. S1C.

### Size exclusion chromatographic separation of intact circulatory IgA assemblies

Size exclusion chromatography (SEC) was used to separate intact IgA assemblies directly in healthy donor plasma/serum for assembly-specific IgA1 Fab clonal profiling. This method was first optimized with SEC fractionation of intact IgA assemblies affinity-purified from healthy human plasma/serum and of total plasma/serum, which were analyzed by mass photometry and bottom-up LC-MS/MS proteomics to validate the separation.

*Size exclusion chromatography conditions.* The same conditions were used for all SEC runs described in this study. A Superdex 200 Increase 10/300 GL column (30 cm length × 10 mm i.d.; 8.6 µm particle size; Cytiva, Marlborough, Massachusetts, USA) was used for all size-exclusion chromatographic separations. All SEC experiments were conducted using an Agilent 1260 Infinity HPLC system (Agilent Technologies, Waldbronn, Germany) with an autosampler, binary pump, column compartment, multi-wavelength detector, and fraction collection module. Both the autosampler and fraction collection modules were chilled to 4 °C to minimize sample degradation, while the column compartment was maintained at 17 °C. Proteins were eluted isocratically within 120 min, with the mobile phase buffer consisting of PBS supplemented with 500 mM NaCl and the flow rate set to 250 µL/min. Absorbance at 280 nm was monitored and recorded for each chromatogram. Fractions were collected every 1 min during the 30-110 min time window, for a total of 80 fractions of 250 µL each.

*SEC method optimization and validation.* The SEC separation of intact IgA assemblies was validated on two samples, for which either 100 µL of plasma/serum from an individual healthy donor diluted 5-fold in PBS (total volume 500 µL) or 100 µL of intact IgA affinity-purified from the same donor plasma/serum sample (stabilized by addition of Tris base as described above) was injected. For whole plasma/serum, fractions were collected the same as described for assembly-specific IgA1 Fab clonal profiling, and then each fraction was subjected to bottom-up sample preparation for analysis by LC-MS/MS proteomics. For affinity-purified IgA, a total of 50 fractions were collected over the time frame of 30-85 min as follows: every 0.5 min (125 µL), 30-50 min; every 4 min (1 mL), 50-62 min; every 2 min (500 µL), 62-70 min; every 5 min (1.25 mL), 70-85 min. These fractions were analyzed by bottom-up proteomics as well as mass photometry.

*Assembly-specific IgA1 Fab clonal profiling.* Individual healthy donor plasma/serum (100 µL) was first diluted 5-fold in PBS, and 500 µL of diluted sample was injected. Fractions were collected every 1 min (250 µL) over the time frame of 30-110 min, for a total of 80 fractions. IgA-containing fractions eluting in the time window of 32-56 min were used for assembly-specific clonal profiling. IgA1 Fab samples were prepared from each SEC fraction as detailed in the following section and subsequently analyzed by intact RPLC-MS.

**IgA1 Fab sample preparation from SEC fractions**

*Affinity purification of SEC-separated intact IgA.* Initial preparation of CaptureSelect IgA Affinity Matrix for affinity purification of all intact IgA from individual fractions following SEC separation of donor serum/plasma was performed as described above (§ Affinity purification of intact circulatory IgA assemblies from individual donor serum/plasma samples). For IgA1 Fab clonal profiling from SEC-fractionated donor plasma/serum, the entire 250-µL fraction volume was added to each prepared spin column with affinity resin together with 200 ng of both recombinant monomeric IgA1 mAbs (7D8-IgA1 and 5D5v2-IgA1) spiked-in as internal standards. Samples were then incubated on an Eppendorf thermal shaker at 750 rpm and room temperature for 1 hr to allow binding of all IgA to the affinity resin beads. After incubation the plugs were removed from the spin columns, and the flow-through was collected via centrifugation for 1 min at 500 × *g*.

*Digestion of IgA1 hinge region.* For IgA1 Fab clonal profiling from SEC-fractionated donor plasma/serum, the beads were washed four times with PBS. Then 50 µL of PBS containing OpeRATOR® and SialEXO® enzymes (Genovis, Kävlinge, Sweden) was added to each affinity-captured IgA sample for overnight (~16 hr) digestion at 37 °C and 750 rpm shaking. The following day, Ni-NTA resin was washed three times with PBS, and 20 µL of prepared resin slurry (1:1 beads:PBS) was added to each sample for immobilization of the His-tagged enzymes. After incubation at room temperature and 750 rpm shaking for 30 min, the enzyme-released IgA1 Fab molecules were collected in the flow-through (final volume ~50-60 µL) via centrifugation for 1 min at 1,000 × *g*. Collected IgA1 Fabs were stored briefly at 4 °C until subjected to intact LC-MS analysis a short time later. For recovery of the IgA Fc molecules/cleavage products still immobilized to the CaptureSelect affinity resin, beads were washed twice more with 150 µL PBS followed by two washes with 150 µL Milli-Q water. Then 100 µL of 0.4% formic acid in Milli-Q water was added for elution of bound IgA Fc molecules from the beads. The IgA1 Fc cleavage products and the intact IgA2 in the supernatant were collected via centrifugation for 1 min at 1,000 × g and stored at −80 °C.

**Intact RPLC-MS profiling of IgA1 Fabs generated from individual SEC fractions**

IgA1 Fabs prepared from individual fractions containing either intact IgA monomer or J-coupled dimer following SEC separation of plasma/serum from Donors 2 and 4 were subjected to intact LC-MS analysis. Fab samples were separated by reversed phase liquid chromatography (RPLC) and analyzed with a Vanquish Flex UHPLC system (ThermoFisher Scientific) coupled to an Orbitrap Exploris 480 mass spectrometer (ThermoFisher Scientific). A MAbPac™ reversed phase analytical column (1 mm i.d. × 150 mm length, polystyrene divinylbenzene stationary phase, 4 µm particle size, 1500 Å pore size) was used for chromatographic separation, and both the column preheater and analytical column ovens were maintained at 80 °C. The injection volume of each Fab sample was 20 µL, which were subsequently separated with a 62-min gradient elution method using mobile phases A (0.1% HCOOH in Milli-Q water) and B (0.1% HCOOH in $CH_3CN$) at a flow rate of 150 µL/min. The initial solvent composition of 10% B was first ramped up to 25% B during the first min (0-1 min), after which the gradient was gradually increased to 40% B in 54 min (1-55 min). Another 1-min ramp to 95% B (55-56 min) before the column was washed by holding at 95% B for 1 min (56-57 min) before returning to the initial 10% B condition in 1 min (57-58 min) which was maintained for the final 4 min (58-62 min). Mass spectra were acquired with the instrument operating in Intact Protein and Low-Pressure mode. MS1 scans were acquired in the range of 500-4,000 m/z with a set resolution of 7,500 (defined at 200 m/z), AGC target of 250%, and a maximum injection time of 50 ms. For each scan, 5 µscans were recorded.

**Data deconvolution and analysis of IgA1 Fab clonal profiles**

Raw files from intact RPLC-MS analysis of IgA1 Fab samples were processed using BioPharmaFinder 3.2 (Thermo Scientific) to determine the intact masses and chromatographic profile of all molecules. Deconvolution of liquid chromatographic-mass spectral data acquired within the 5-57 min timeframe of each raw file was performed using 0.1 min sliding windows with a 25% offset with the ReSpect algorithm. Merge tolerance and noise rejection parameters were set at 30 ppm and 95%, respectively. Deconvolution output masses were bounded by the range of 10,000 to 100,000, and 48,000 was defined as the target mass. The Intact Protein peak model was selected for mass deconvolution, with charge states between 10 and 60 included and the mass tolerance set to 30 ppm.

As previously described (***6, 7***), BioPharmaFinder component identifications were further processed and analyzed using Python 3.8.10 (***8***) (with an in-house python library in addition to free software libraries: Pandas 1.5.0, Numpy 1.23.3 (***9***), Scipy 1.9.1 (***10***), Matplotlib 3.6.0 (***11***), Seaborn 0.12.0). Data were filtered to remove non-Fab identifications, with only those components meeting all criteria (mass of 45-53 kDa, most intense charge state >1000 m/z, score of ≥40) considered as IgA1 Fab molecules. Masses were recalculated using an intensity-weighted mean with only the most intense peaks representing the top 90% of the total intensity included. Intensities were normalized to the 7D8-IgA1 and 5D5v2-IgA1 mAb standards, with relative mass and retention time shifts applied to minimize mass error and RT deviation between runs. After removal of the mAb Fabs from the data, endogenous clones were matched between runs using average linkage (unweighted pair group method with arithmetic mean UPGMA) $L_\infty$ distance hierarchical clustering. Flat clusters were formed based on a cophenetic distance constraint derived from the mass and retention time tolerances set to 2 Da and 1 min, respectively. Clones within a flat cluster were considered identical between runs. Hierarchical clustering of samples was performed using correlation distance and UPGMA average linkage, visualization of the tree was made using iTOL.

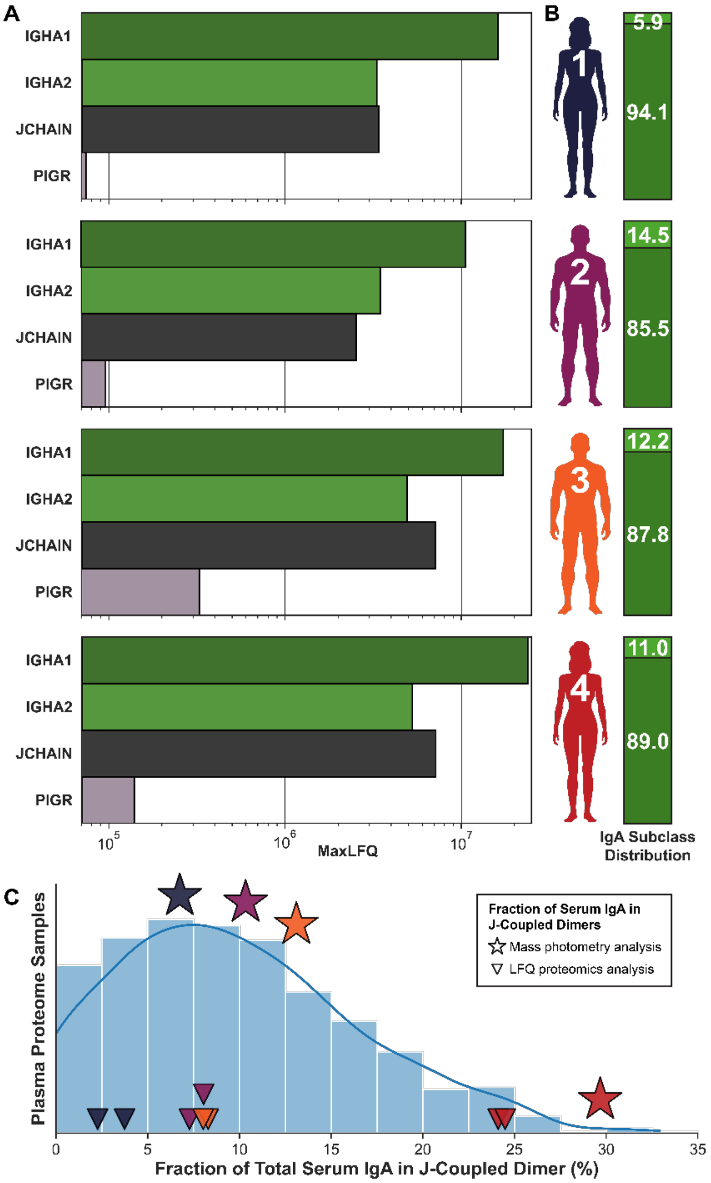


**Fig. S1. Bottom-up proteomics confirms each donor's distribution of circulatory IgA assemblies.**

**(A)** MS-based bottom-up proteomics analysis of circulatory IgA assemblies affinity-purified from Donors 1-4 as indicated, with the abundances of IGHA1, IGHA2, JCHAIN, and PIGR shown. The lack of PIGR (>100-fold lower than IGHA) confirms no SIgA or larger assemblies are present in the distribution in Fig. 1B.

**(B)** Each bar chart depicts the IgA subclass distribution (IgA1, dark green; IgA2, light green) determined from MS-based bottom-up proteomics analysis of Donor 1-4 serum/plasma.

**(C)** Histogram depicting the circulatory IgA assembly distribution (expressed as the fractional abundance of J-coupled dimer) calculated for a publicly-available plasma proteomics dataset (N = 687 sampling points, 139 patients; see Supplemental Materials and Methods). Annotated symbols correspond to IgA assembly distribution for Donors 1-4 observed by mass photometry (stars) or calculated from bottom-up proteomics analysis of serum/plasma using two related approaches to correct for the presence of J-coupled IgM pentameric assemblies in circulation (triangles). Stars and triangles are color-coded according to the corresponding color of Donors 1-4 as shown in the above panel.

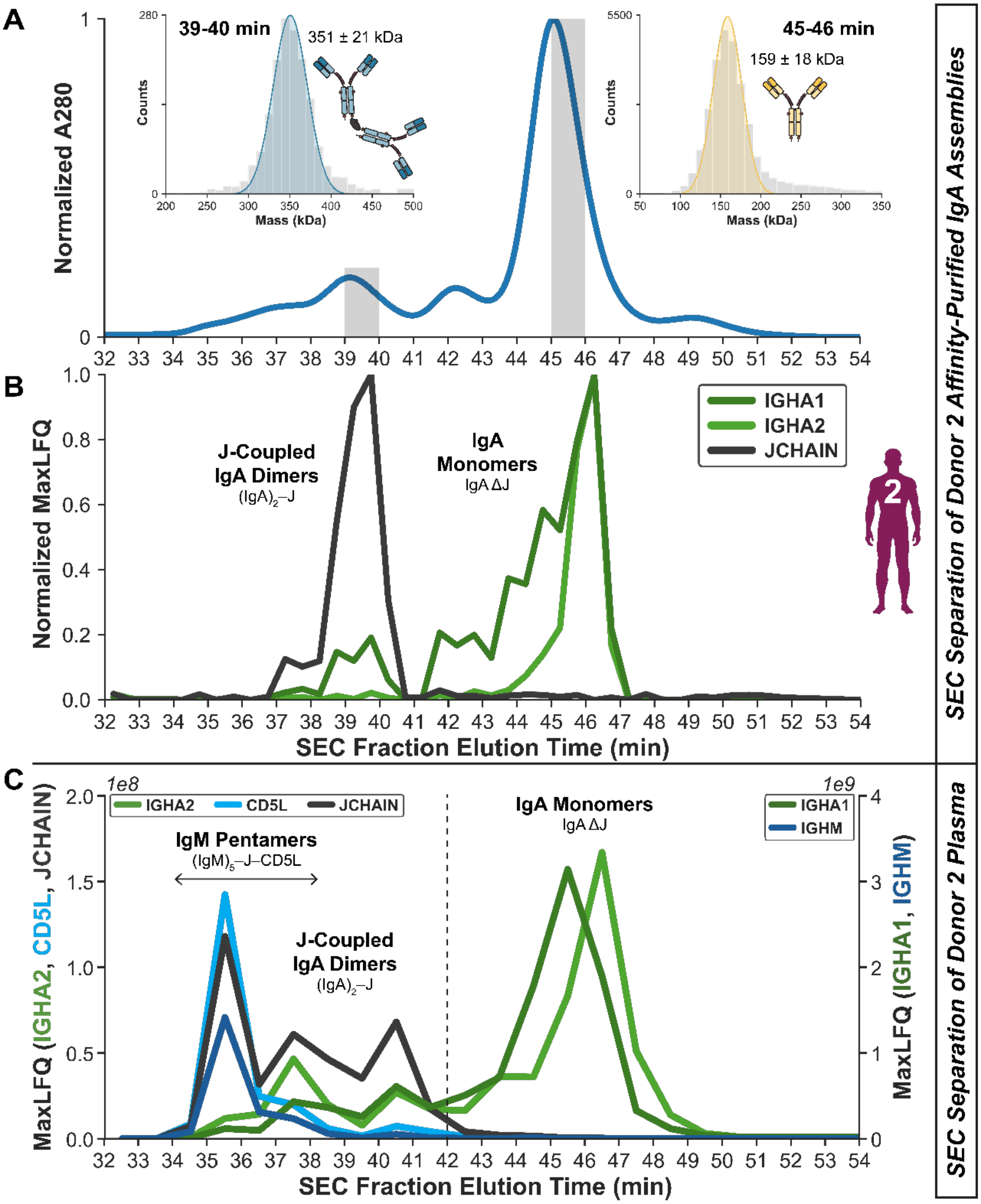


**Fig. S2. Size exclusion chromatographic separation of J-coupled dimers and monomers of IgA.**

**(A)** Size exclusion chromatogram of affinity-purified circulatory IgA assemblies of Donor 2, depicting the absorbance at 280 nm as a function of time. The two inset panels depict the mass distribution of the indicated fractions (also highlighted in grey) determined by single-molecule mass photometry.

**(B)** Results from MS-based bottom-up proteomics analysis of each SEC fraction from **A**.

**(C)** Results from MS-based bottom-up proteomics analysis of SEC-fractionated plasma of Donor 2.

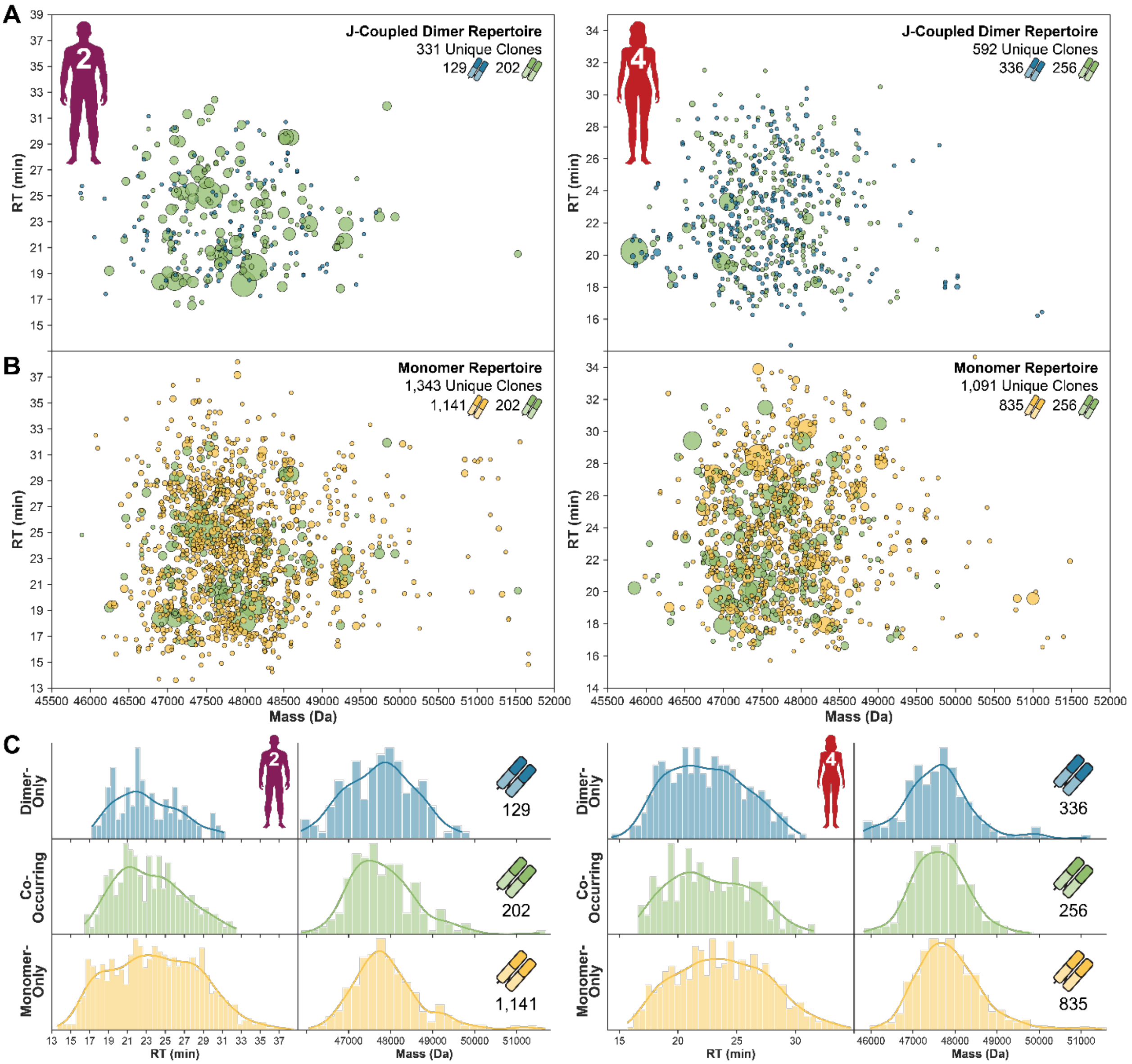


**Fig. S3. Monomer and J-coupled dimer repertoires each contain hundreds of unique IgA1 clones.**

**(A)** All unique IgA1 Fab clones detected from J-coupled dimers for Donor 2 **(left)** and Donor 4 **(right)**. Each unique clone is plotted according to its characteristic mass and RPLC RT. The relative abundance of each is indicated by the scaling of the marker size. The total number of unique clones, including a breakdown of each of the three types, is annotated at top right. The color scheme used is established in the Main Text, with dimer-only clones in blue, co-occurring clones in green, and monomer-only clones in yellow.

**(B)** Same as in **A** except clones shown were detected from monomers.

**(C)** Histograms depicting the distribution of characteristic RPLC RT **(left)** and mass **(right)** identifiers for each unique IgA1 Fab clone, which are separated according to the three unique clone types as indicated.

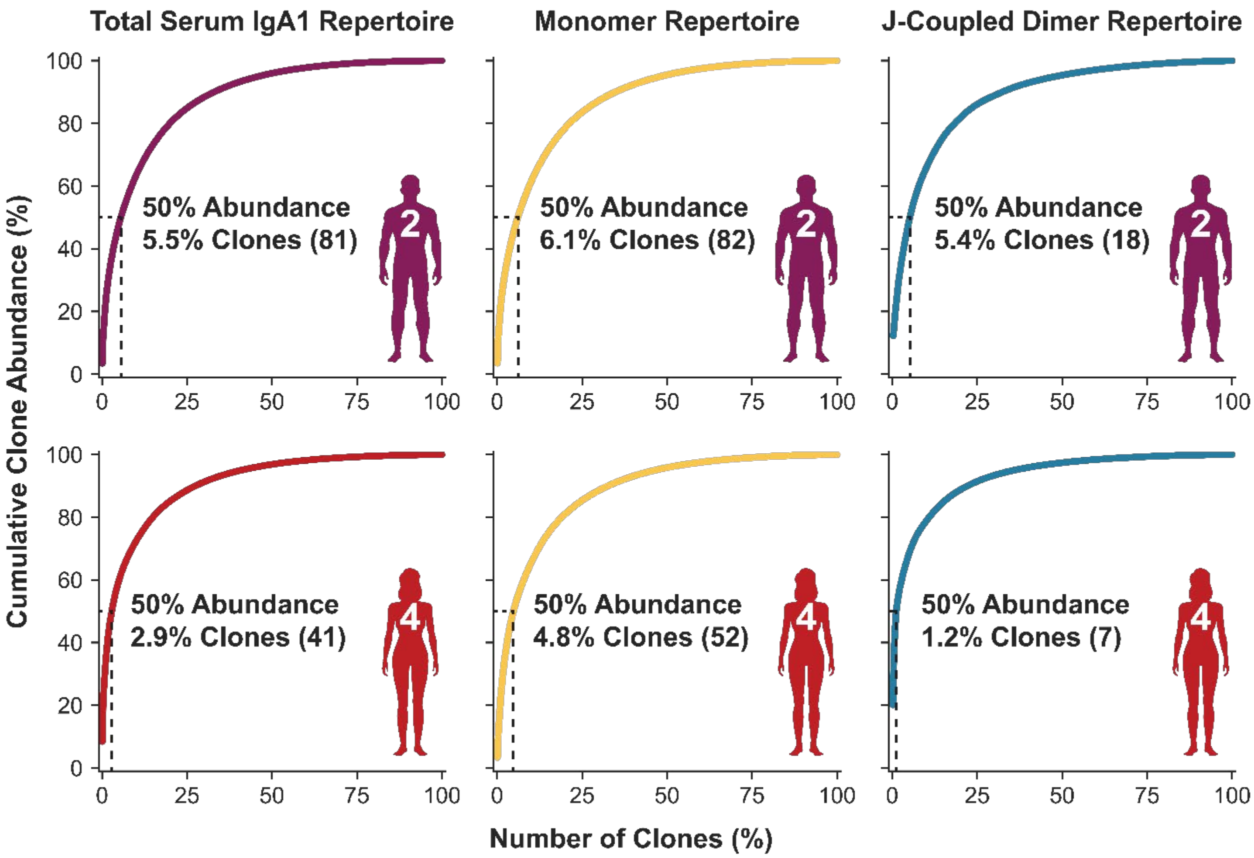


**Fig. S4. Serum IgA1 clonal repertoires are simple.**

Cumulative clone abundance as a function of the proportion of clones in the total **(left)**, monomer **(middle)**, and J-coupled dimer **(right)** repertoires of Donor 2 **(top)** and Donor 4 **(bottom)**. The dashed vertical and horizontal lines in each panel indicate the proportion of clones (labeled in each, with the equivalent number of unique clone counts shown in parentheses) which account for 50% of the cumulative abundance in each repertoire.

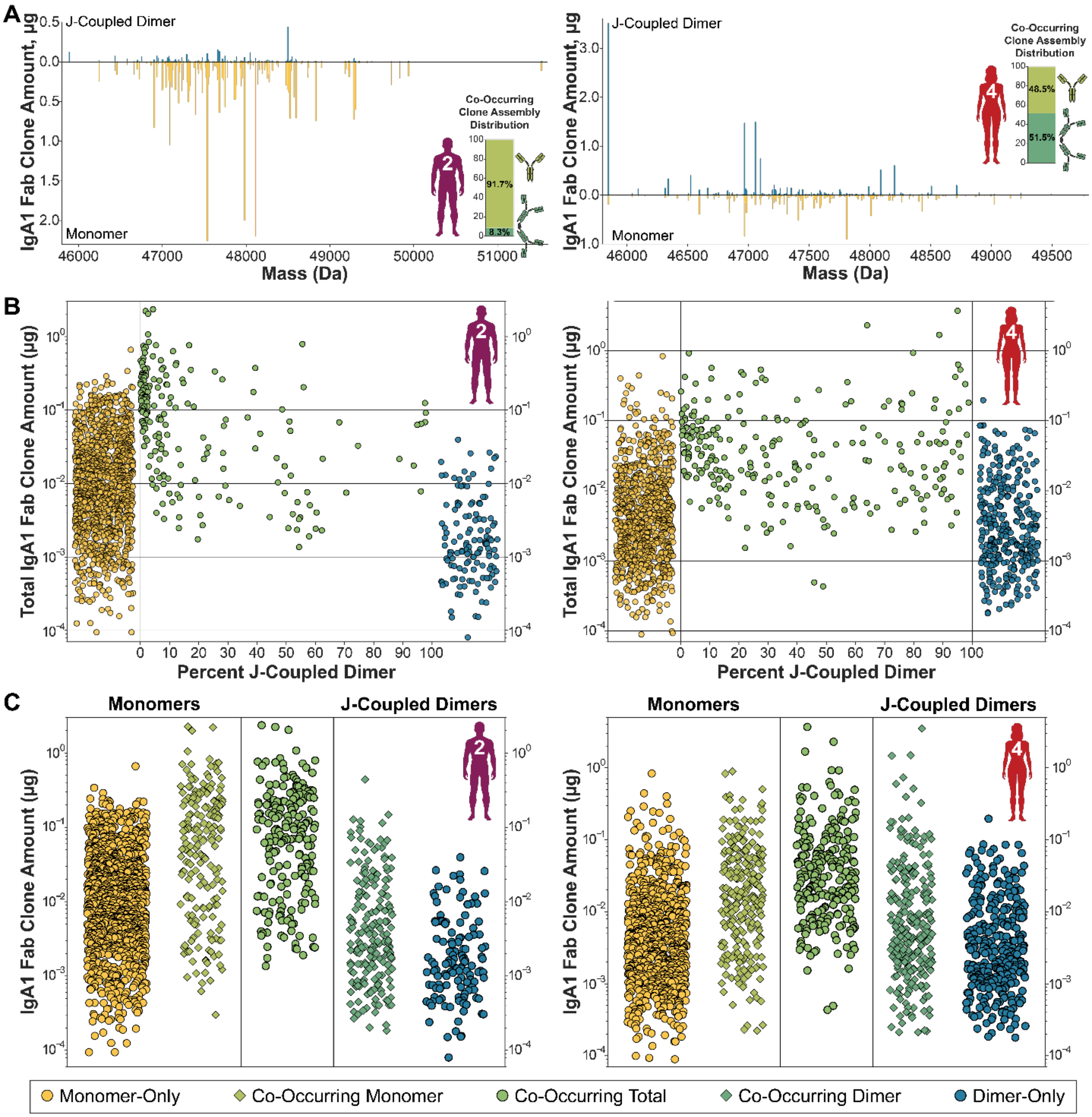


**Fig. S5. The distribution of co-occurring clones differs between donors.**

**(A)** Profile of all co-occurring IgA1 clones for Donor 2 **(left)** and Donor 4 **(right)**. Each stick represents a unique co-occurring clone, plotted according to its characteristic mass. The total abundance of each is divided according to the fractional contributions of the J-coupled dimer (blue, upper panel) and monomer (yellow, lower panel) forms. Inset bar charts indicate the cumulative contributions of the monomer (yellow-green) and J-coupled dimer (darker blue-green) abundances of all co-occurring clones.

**(B)** Distribution of all unique clones according to abundance and assembly distribution.

**(C)** Similar to **B** but with the assembly-specific amount of each unique co-occurring clone shown and with clones grouped by category as indicated.

**Table S1.** Healthy donor sample details.

| *Donor ID* | *Sex* | *Age* | *Sample Details* | *Sample Source* |
|---|---|---|---|---|
| ***Donor 1*** | Female | 58 | Serum | Sanquin Research |
| ***Donor 2*** | Male | 48 | $K_2$EDTA Plasma | Zen-Bio Inc |
| ***Donor 3*** | Male | 60 | Serum | Sanquin Research |
| ***Donor 4*** | Female | 60 | Serum | Sanquin Research |

**Table S2.** Summary of results from analysis of affinity-purified circulatory intact IgA assemblies from four individual healthy donors by single-molecule mass photometry.

| | | Donor 1 | Donor 2 | Donor 3 | Donor 4 |
|---|---|---|---|---|---|
| **Mass (kDa, μ ± σ)** | Monomer | 155 ± 15 | 160 ± 15 | 153 ± 15 | 157 ± 15 |
| | J-coupled dimer | 335 ± 21 | 343 ± 24 | 335 ± 21 | 343 ± 20 |
| **Fraction of all intact IgA assemblies detected** | Monomer | 96.5 % | 94.5 % | 93.0 % | 82.6 % |
| | J-coupled dimer | 3.5 % | 5.5 % | 7.0 % | 17.4 % |
| **Fraction of total serum IgA** | Monomer | 93.2 % | 89.7 % | 86.9 % | 70.3 % |
| | J-coupled dimer | 6.8 % | 10.3 % | 13.1 % | 29.7 % |

**Table S3.** Summary of results from assembly-specific LC-MS profiling analysis of endogenous circulatory IgA1 clonal repertoires.

### Donor 2 Circulatory IgA1 Clonal Profiling Results

| Population | Number of Unique Clones | Total Serum IgA1 Repertoire | Monomer Repertoire | J-Coupled Dimer Repertoire |
|---|---|---|---|---|
| Monomer-Only | 1141 | 77.5% | 85.0% | |
| Co-Occurring | 202 | 13.7% | 15.0% | 61.0% |
| Dimer-Only | 129 | 8.8% | | 39.0% |
| **All Clones** | **1472** | **100%** | | |
| **All Monomers** | **1343** | 91.2% | **100%** | |
| **All Dimers** | **331** | 22.5% | | **100%** |

| Population | Cumulative Clone Amount (µg) | Total Serum IgA1 Repertoire | Monomer Repertoire | J-Coupled Dimer Repertoire | Co-Occurring Repertoire |
|---|---|---|---|---|---|
| Monomer-Only | 30.393 | 44.5% | 47.0% | | |
| Co-Occurring Monomer | 34.313 | 50.2% | 53.0% | | 91.7% |
| Co-Occurring Dimer | 3.105 | 4.5% | | 86.1% | 8.3% |
| Dimer-Only | 0.502 | 0.7% | | 13.9% | |
| **All Clones** | **68.312** | **100%** | | | |
| **All Monomers** | **64.705** | 94.7% | **100%** | | |
| **All Dimers** | **3.607** | 5.3% | | **100%** | |
| **All Co-Occurring** | **37.418** | 54.8% | | | **100%** |

### Donor 4 Circulatory IgA1 Clonal Profiling Results

| Population | Number of Unique Clones | Total Serum IgA1 Repertoire | Monomer Repertoire | J-Coupled Dimer Repertoire |
|---|---|---|---|---|
| Monomer-Only | 835 | 58.5% | 76.5% | |
| Co-Occurring | 256 | 17.9% | 23.5% | 43.2% |
| Dimer-Only | 336 | 23.5% | | 56.8% |
| **All Clones** | **1427** | **100%** | | |
| **All Monomers** | **1091** | 76.5% | **100%** | |
| **All Dimers** | **592** | 41.5% | | **100%** |

| Population | Cumulative Clone Amount (µg) | Total Serum IgA1 Repertoire | Monomer Repertoire | J-Coupled Dimer Repertoire | Co-Occurring Repertoire |
|---|---|---|---|---|---|
| Monomer-Only | 12.775 | 29.2% | 48.6% | | |
| Co-Occurring Monomer | 13.506 | 30.9% | 51.4% | | 48.5% |
| Co-Occurring Dimer | 14.314 | 32.8% | | 82.2% | 51.5% |
| Dimer-Only | 3.101 | 7.1% | | 17.8% | |
| **All Clones** | **43.697** | **100%** | | | |
| **All Monomers** | **26.281** | 60.1% | **100%** | | |
| **All Dimers** | **17.415** | 39.9% | | **100%** | |
| **All Co-Occurring** | **27.820** | 63.7% | | | **100%** |

### SI References


1. F. Soltermann *et al.*, Quantifying Protein–Protein Interactions by Molecular Counting with Mass Photometry. *Angew. Chem. Int. Ed. Engl.* **59**, 10774-10779 (2020).
2. P. Skowronek *et al.*, Rapid and In-Depth Coverage of the (Phospho-)Proteome With Deep Libraries and Optimal Window Design for dia-PASEF. *Mol. Cell. Proteomics* **21**, 100279 (2022).
3. V. Demichev, C. B. Messner, S. I. Vernardis, K. S. Lilley, M. Ralser, DIA-NN: neural networks and interference correction enable deep proteome coverage in high throughput. *Nat. Methods* **17**, 41-44 (2020).
4. M.-A. Gerault, L. Camoin, S. Granjeaud, DIAgui: a Shiny application to process the output from DIA-NN. *Bioinform. Adv.* **4**, vbae001 (2024).
5. V. Demichev *et al.*, A time-resolved proteomic and prognostic map of COVID-19. *Cell Syst.* **12**, 780-794.e787 (2021).
6. A. Bondt, K. A. Dingess, M. Hoek, D. M. H. van Rijswijck, A. J. R. Heck, A Direct MS-Based Approach to Profile Human Milk Secretory Immunoglobulin A (IgA1) Reveals Donor-Specific Clonal Repertoires With High Longitudinal Stability. *Front. Immunol.* **12**, 789748 (2021).
7. K. A. Dingess *et al.*, Identification of common and distinct origins of human serum and breastmilk IgA1 by mass spectrometry-based clonal profiling. *Cell. Mol. Immunol.* **20**, 26-37 (2023).
8. W. McKinney, Data structures for statistical computing in Python. *Proceedings of the 9th Python in Science Conference* 56-61 (2010).
9. S. v. d. Walt, S. C. Colbert, G. Varoquaux, The NumPy Array: A Structure for Efficient Numerical Computation. *Comput. Sci. Eng.* **13**, 22-30 (2011).
10. P. Virtanen *et al.*, SciPy 1.0: fundamental algorithms for scientific computing in Python. *Nat. Methods* **17**, 261-272 (2020).
11. J. D. Hunter, Matplotlib: A 2D Graphics Environment. *Comput. Sci. Eng.* **9**, 90-95 (2007).
